# A Population Vector Model of Visual Working Memory for Real-World Scenes

**DOI:** 10.64898/2026.01.23.701256

**Authors:** John E. Kiat, Steven J. Luck

## Abstract

Visual working memory is essential for navigating through and interacting with complex real-world environments. It is therefore important to understand how natural visual inputs—characterized by complex contours, continuously varying feature gradients, and spatial relationships—are represented in working memory. However, most research in this field has focused on simplified arrays of discrete artificial objects, favoring experimental control and modeling simplicity over ecological validity. This has led to quantitative models of working memory that require inputs consisting of easily parsed objects defined by a single value along one or more simple feature dimensions. It is not clear how these models could be updated to represent complex, photograph-like scenes. To overcome this limitation, we introduce a *population vector model of working memory* that was designed specifically for real-world scenes. This model represents a scene as a noisy vector of neural firing rates across one or more areas of the ventral pathway, as estimated by a deep neural network model. We show that this model can account for both variations in behavioral performance and patterns of brain activity in tasks that require storing naturalistic scenes in working memory. These results demonstrate the viability of our general modeling approach, setting the stage for more sophisticated models that can fully account for the storage of real-world scenes in working memory.

**Public Significance Statement:** People unconsciously store visual information briefly in memory thousands of times each day and use this information to help them perform a broad range of natural tasks. Although the visual working memory system used for this purpose has been intensively studied using simple and highly controlled experimental stimuli (e.g., arrays of colored squares), there has been little progress in developing formal quantitative accounts of how real-world scenes are stored in this system. Here, we provide a new model of visual working memory that was designed for real-world scenes and can predict both behavior and brain activity when people store these scenes in memory.

---

Visual working memory is a fundamental cognitive system that stores information temporarily to serve the needs of ongoing tasks (Bays et al., 2024; Luck & Vogel, 2013; Schurgin, 2018). This system holds representations of what we are seeking when we search the world for objects (Carlisle et al., 2011; Gunseli et al., 2014). It is the representational system used to represent objects undergoing mental rotation (Hyun & Luck, 2007). It is used perhaps 100,000 times each day in the context of eye movements, allowing our visual systems to compare the object we planned to fixate with the object we actually fixated (Hollingworth et al., 2008; Hollingworth & Luck, 2009). Furthermore, it has become a key testbed for studying the nature and neural storage of mental representations (Barbosa et al., 2021; Funahashi et al., 1989; Lundqvist et al., 2016; Todd & Marois, 2004), for exploring the limitations of human cognition (Oberauer, 2019; Saults & Cowan, 2007; Vogel & Awh, 2008), and for understanding why people differ in intellectual ability (Fukuda et al., 2010; Unsworth & Engle, 2007).

Although psychologists and neuroscientists have studied visual working memory since at least the 1960s, research in this area started expanding rapidly in the mid-1990s with the popularization of two distinct experimental approaches. In one approach, the observer sees a photograph of a real-world scene, followed by a brief delay or some other kind of disruption, and then an altered version of the scene (Grimes, 1996; O’Regan et al., 2000; Rensink et al., 1997). These two versions alternate until the observer is able to determine whether the two versions differ. The central finding from research using this approach is that many alternations are typically needed for a person to detect a fairly substantial difference between the two scenes. This has been termed *change blindness*, and it has long caught the interest of both philosophers and psychologists because it contrasts with the richly detailed phenomenological experience people have while each scene is visible (Dretske, 2007; Noe, 2002; Scholl & Simons, 2001).

In the other main approach (Jiang et al., 2000; Luck & Vogel, 1997; Pashler, 1988; Phillips, 1974), the observer briefly views an array of discrete, simple, artificial objects (e.g., six colored squares), followed by a delay interval and then a new array that is either identical or differs in the properties of one object (e.g., a change in the color of one of the six squares). The observer reports whether or not a change was detected. The central finding from research using this approach is that change detection accuracy is near ceiling for arrays containing one or two objects (assuming that the changes are large) but decreases rapidly as the number of items in the arrays (the *set size*) is increased (Luck & Vogel, 2013). This pattern of results indicates that the storage capacity of visual working memory is quite small. Later studies modified this task to require that observers report exact feature values instead of detecting changes, and the precision of these reports also declines as the number of items increases (Adam et al., 2017; Bays, 2014; Nassar et al., 2018; Schurgin et al., 2020; Wilken & Ma, 2004; W. Zhang & Luck, 2008).

Studies using arrays of simple artificial objects have come to dominate the field, and research on visual working memory using natural scenes has become relatively rare. The use of simple, discrete objects has been valuable because this approach makes it easy to manipulate well-understood stimulus properties and to develop formal models of task performance. The resulting models require that the visual input has been parsed into a set of discrete items, each of which has a single feature value along one or more continuous dimensions (Bays, 2014; Rouder et al., 2008; Schurgin et al., 2020; van den Berg et al., 2014; W. Zhang & Luck, 2008).

## The Challenge of Representing Real-World Scenes in Working Memory

Although useful for testing hypotheses about the nature of working memory, item-based models currently cannot be applied to real-world scenes like the one shown in Figure 1. For example, these models have no means of capturing the fact that the surface of an object may vary gradually in hue or brightness (like the sky in Figure 1), no means of representing complex shape contours (like the hills), no means of representing spatial relationships (like the spacing of the boulders), and no means of representing scene regions that cannot be parsed into discrete and homogeneous items (like the clouds). There is currently no clear path forward that would eventually allow these models to capture the key elements of real-world scenes.

**Figure 1.**
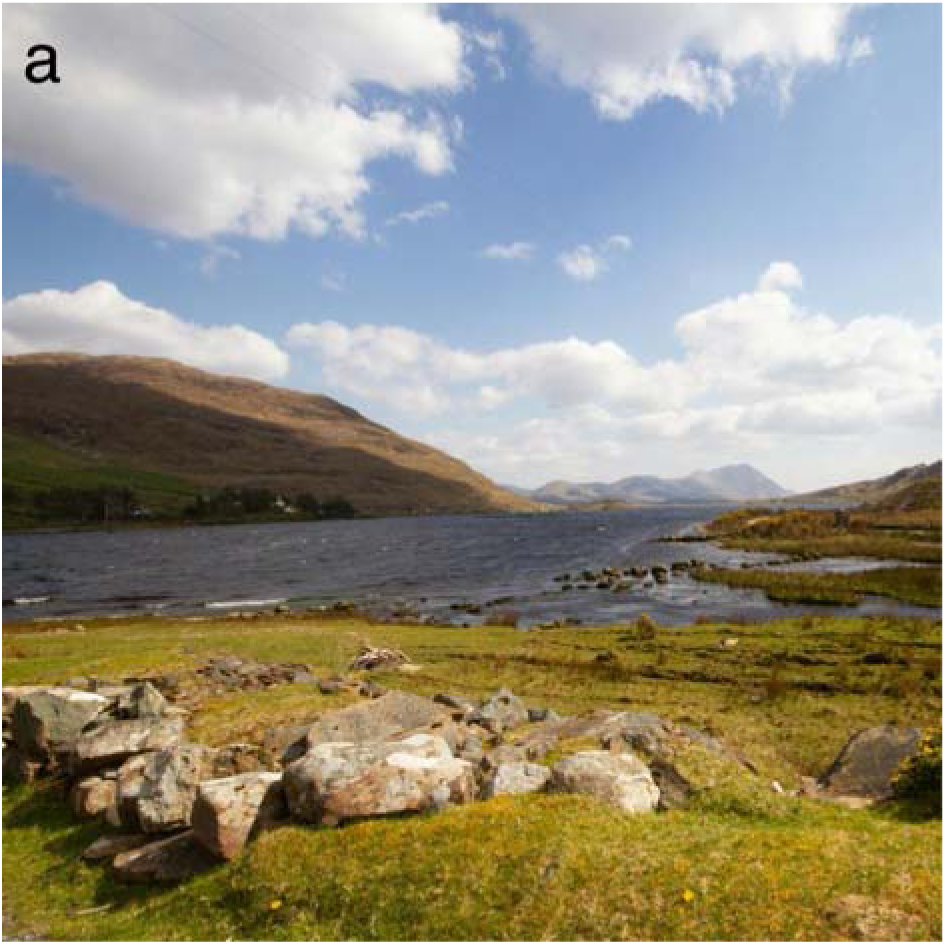
Example of the type of scene that the present model is designed to handle. *Note*. Scenes like this cannot readily be represented by models designed for arrays of discrete, simple, artificial objects. For example, the color of the sky is not a single value but changes gradually over space, and the contours of the hills vary continuously rather than containing a single orientation.

Although it might seem as if the scene in Figure 1 could be parsed into a set of discrete items, with each item containing one feature value per dimension, this is far from straightforward. For example, suppose that the cloud in the upper left corner is taken to be a single item. In that framing, it is unclear how the gradually changing luminance across the surface of the cloud could be represented. If we instead treat the cloud as a set of multiple overlapping items, each with relatively homogenous shading captured in a single luminance value, it is unclear how many regions are required and how the relationships between them are stored to represent the overall luminance gradient.

Moreover, it is not clear what features would be used to represent the amorphous shape of the cloud. Even in simpler cases, shape is notoriously difficult to represent in terms of simple continuous features. There are some approaches for this, such as Fourier descriptors (Zahn & Roskies, 1972). However, these approaches still require storing multiple interrelated values to represent a single shape (violating the assumption of one feature value per dimension). Indeed, machine vision research has almost entirely moved away from using an alphabet of simple features to represent objects, opting instead for neural network and transformer approaches (Abbas et al., 2019; Tsourounis et al., 2022).

Furthermore, models of working memory based on independent items have no way of encoding the relative sizes and spatial positions of objects, like the positions and sizes of the mountains relative to the boulders in Figure 1. Indeed, Clevenger & Hummel (2014) attempted to create an item-based model that included spatial and size relationships for simple artificial objects, and it was extremely challenging to account for all possible pairs of relationships. Thus, there is currently no clear path forward that would eventually allow these item-based models to capture the key elements of real-world scenes.

This does not mean that item-based working memory representations play no role in real-world visual tasks. For example, if observers were asked to count the number of boulders given a brief presentation of the image in Figure 1, they could perform this task by storing a single “pointer” to each boulder in working memory. In addition, visual working memory has been proposed to play an essential role in gaze control that could be achieved with simple item-based representations (Hollingworth et al., 2008; Hollingworth & Luck, 2009). However, a different kind of working memory representation seems to be necessary to capture the key properties of an entire scene, one that can handle color gradients, complex shapes, relational information, and other important elements of real-world scenes.

## Overview of the Present Approach

Here, we introduce an approach that is designed to quantitatively model the working memory representations of arbitrarily complex naturalistic scenes. Our approach begins by assuming that—to a first approximation—the working memory representation of a scene is simply a weak, noisy version of the perceptual representation of that scene in one or more regions of the ventral visual processing stream. This assumption is supported by studies of single-unit activity in inferotemporal (IT) cortex, in which a neuron’s firing rate during the delay period of a working memory task is approximately proportional to the firing rate measured during the encoding period (Chelazzi et al., 1998; Woloszyn & Sheinberg, 2009). We therefore conceive of the working memory representation of a scene as a weaker version of the brain’s perceptual representation of a scene in one or more areas of the ventral pathway. This weakened activation pattern is closer to the background firing rates of the neurons and, therefore, functionally noisier.

Because there is no method currently capable of measuring the pattern of single-unit firing rates across the entire human ventral visual processing pathway, we currently use a deep neural network (DNN) model of the ventral stream called CORnet (Kubilius et al., 2018) to model the brain’s perceptual representation of a given scene. CORnet was explicitly structured to reflect several key properties of the ventral stream. For example, as illustrated in Figure 2a, CORnet contains separate modules corresponding to the main visual areas of the ventral stream (V1, V2, V4, and IT), which vary in the size of the receptive fields and the abstractness of the features. Each CORnet area consists of multiple layers of units, just as cortical visual areas contain multiple layers of neurons. The specific version of CORnet used here (*CORnet-S*) also simulates within-area recurrent connections. CORnet-S scores well on the *Brain-Score* metric (Schrimpf et al., 2020) of similarity to measurements of neural activity in the ventral stream.

**Figure 2.**
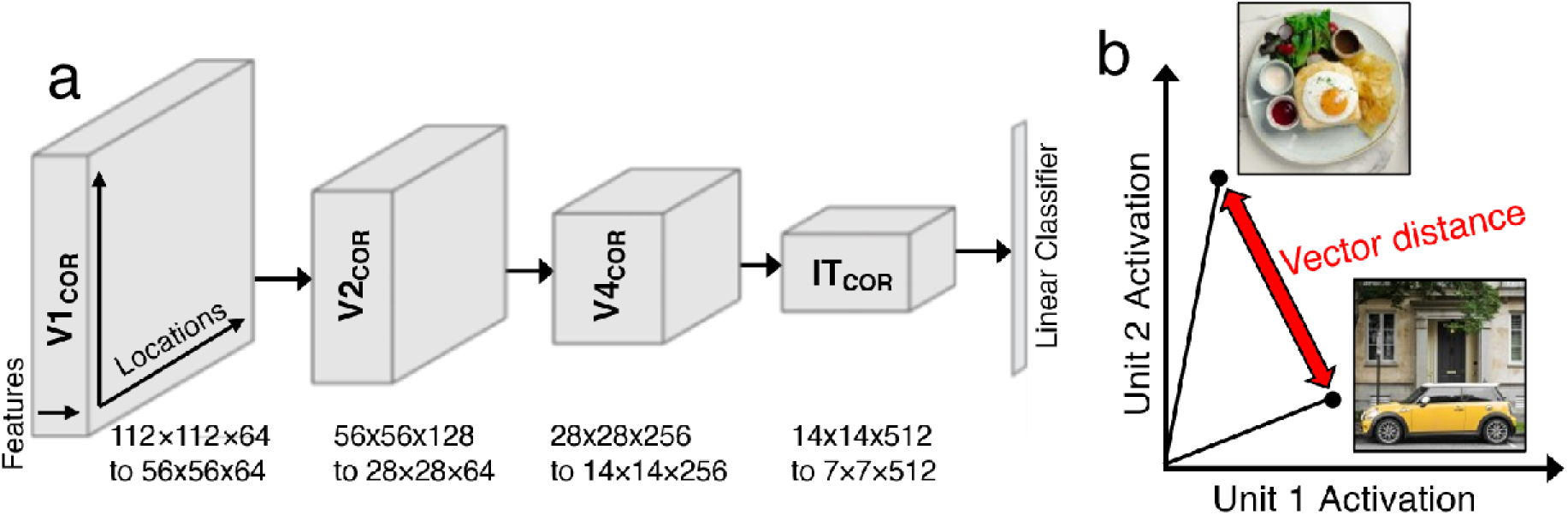
Essence of the population vector model. *Note*. **a**, CORnet-S model of scene processing, which is used to form an activation vector for each scene. It consists of four areas (V1_COR_, V2_COR_, V4_COR_, IT_COR_), each composed of multiple layers. The input is a 2D color bitmap of a scene, which creates a pattern of activation over the units in the input layer of V1_COR_. The activation passes through weights from one layer to the next and from one area to the next, creating a new pattern of activation in each layer. A linear classifier (not used in the present study) uses the pattern of activation in the output layer of IT_COR_ to categorize the scene. The weights are set by training the model to categorize a large set of scenes. **b**, Sketch of how change detection performance is predicted by a simplified version of the model in which a scene is represented by only two units. Any given scene produces an activation value in each of the two units, and the representation of a scene can be considered a vector in this 2D space. The ability of an observer to detect a difference between two scenes is proportional to the distance between the vectors for the two scenes (after the application of a normalization process inspired by Weber’s law). In the full model, the representational space has a huge number of dimensions, one for each of the thousands of units in a given layer of CORnet.

Like DNNs that were developed for engineering applications (Dhillon & Verma, 2020; K. He et al., 2016; Krizhevsky et al., 2012), CORnet takes a two-dimensional array of RGB (red-green-blue) values as its input, and from this input array produces a pattern of activation across simplified neuron-like units. The weights of the connections between units were set by training the model to classify a massive set of labeled photographs of real-world objects (Russakovsky et al., 2015). However, unlike most DNNs, CORnet was not developed to solve engineering problems, but was instead designed for the express purpose of matching the neural representations of scenes across the major stages of the ventral visual processing pathway. We therefore use the pattern of activation across units in CORnet for a given scene as an approximation of the perceptual representation of that scene, with a weakened version of this pattern serving as the working memory representation of the scene.

The pattern of activation across a set of units in a DNN can be conceived as a vector in a massively multidimensional space (with one dimension for each unit). We therefore call our model the *population vector model of working memory* because it represents a given scene as a weak/noisy version of the vector of activations produced by that scene across the population of neuron-like units. It is important to note that DNNs like CORnet have some well-documented limitations (Baker et al., 2018; Ballester & Araujo, 2016; Gatys et al., 2017; Jozwik et al., 2018; Xu & Vaziri-Pashkam, 2021), which will be addressed in the General Discussion. Our broader theory is that the visual working memory representation of a natural scene can be effectively modeled as a weak/noisy version of the vector of activation across neurons in the ventral stream. Consequently, we are using CORnet here as a practical means of obtaining an approximation of this vector. We are not committed to CORnet per se, but it is a reasonable placeholder until better models of the ventral stream become widely available. Thus, the present study should not be construed as a test of CORnet, but as a test of the viability of a broader theory of visual working memory for naturalistic scenes.

Figure 2b provides a sketch of how our approach links the pattern of CORnet activation to change-detection performance, focusing on the activation values from just two units. Each input scene produces an activation level in each of the two units, and the pattern across the two units can be considered a location (a *vector*) in a 2D space. The distance between the two vectors quantifies the (dis)similarity between the scenes. Each layer of CORnet contains thousands of units, so the distance between vectors ends up being computed from a massively multidimensional space. We normalize the vector distances in a manner inspired by Weber’s law, which specifies that the difference between two sensory inputs needed to reliably discriminate between them scales with the magnitude of the inputs (see Method). This could potentially lead to a form of capacity limitations, because complex scenes will activate more units than simple scenes, and a larger absolute difference between scenes will therefore be necessary to detect a change in complex scenes than in simple scenes. However, our model does not parse the scene into discrete objects, and it lacks an explicit mechanism for producing capacity limitations. Thus, although it may be inevitable that visual working memories are stored as vectors of firing rates, there is a broad range of potential alternative models in which the relevant vectors arise after the scene has been parsed into discrete objects (as in item-based models), after some kind of capacity limit or compression has been applied (as in Bates & Jacobs, 2020; Jakob & Gershman, 2023), after top-down signals have been integrated into the vectors (as in Y. Xie et al., 2023), and so on.

Prior work has already demonstrated the plausibility of using DNN activation vectors to predict working memory performance (Bates et al., 2024), using data from an experiment in which observers were shown a naturalistic scene and then asked to judge which of many scenes was most similar to the original scene (Son et al., 2022). The observed judgments were predicted reasonably well by several different DNNs, indicating that DNNs can capture how humans judge the similarity between a remembered scene and a visible scene. Brady & Störmer (2024) obtained similar results for arrays of isolated objects. The DNNs used in these previous studies were selected for analysis because they perform image classification tasks accurately, not because the pattern of activation across units matches the perceptual representations of the ventral stream. Thus, these studies are not directly relevant for testing our general theory, which is that working memory representations are weak/noisy copies of the perceptual representations within one or more regions of the ventral stream. By contrast, the CORnet DNN used in the present study was expressly designed to be a model of the human ventral stream.

## Overview of Experiments

The primary goal of the present study was to assess the viability of our general theory, in which natural scenes are stored in working memory as weak/noisy copies of the neural activation vectors produced in the ventral stream by the perception of those scenes (using CORnet because it is the best current method of estimating those activation vectors). Because there are no other full-fledged models of working memory for natural scenes, we could not compare our model with other models. We therefore took the approach of determining whether our model is viable and worthy of further development by assessing whether the model could explain a substantial amount of variance in change detection behavior (relative to clear-cut reference points described below) and whether it could account for patterns of brain activity during the delay period. Because all of our data and code are freely available at https://osf.io/7ukgj/, researchers who develop new models can readily compare the performance of these new models with the performance of the present model.

The first three experiments assessed the ability of our model to account for behavioral performance in change detection tasks. Experiment 1 used a *flicker* paradigm like those originally used to examine visual working memory for natural scenes (Grimes, 1996; O’Regan et al., 2000; Rensink et al., 1997), with two versions of a naturalistic scene that alternate until the observer detects the difference between them. Experiment 2 used the same scenes but with a one-shot paradigm like those originally used to examine visual working memory for arrays of simple artificial objects (Jiang et al., 2000; Luck & Vogel, 1997; Pashler, 1988; Phillips, 1974). In this task, the first and second versions of a scene are each shown once, and then the observer reports whether they were the same or different. Experiment 3 used the flicker task of Experiment 1, but with a different set of scene pairs. In all of these experiments, we asked whether the distance between the vectors for the two scenes in a pair could predict how quickly or accurately observers could detect whether the two scenes were different.

We compared the predictive ability of our model with two other sources of predictions that could serve as reference points^1^. First, we considered a lower reference point in which the vector distances were obtained from the set of raw RGB values in the image bitmaps. By definition, these bitmaps contain all the information that is needed to perform the change detection task, and it is therefore possible that they are sufficient to approximate the representational format of working memory. Thus, if the vectors in our model could not substantially outperform the reference point provided by vectors obtained from the RGB representations, this would be evidence against the viability of our theory. Second, we considered an upper reference point in which we asked human observers to judge the perceptual similarity of the scene pairs that were shown side by side for an unlimited amount of time. These explicit ratings of similarity for the image pairs should do an excellent job of predicting how easily observers can determine whether the two images in a pair are the same or different in our change detection tasks. If the amount of variation in change detection performance across scene pairs explained by our model comes anywhere near the amount of variation explained by these ratings of perceptual similarity, then this would indicate that our model has some value.

It might seem as if the ratings of perceptual similarity when the scenes in a pair are viewed side by side would perfectly predict the speed and accuracy of change detection when those same scene pairs are presented sequentially. Indeed, that is why the similarity ratings provide a reasonable reference point that our model should aim for. In reality, however, the similarity ratings might not fully account for change detection performance, either because of imperfections in the ratings or because observers might weight the various image features differently when making an explicit judgment of similarity than when making a same-different judgment. It is therefore possible that our model might explain unique variance in change detection performance after controlling for the explicit similarity ratings. This would provide even greater reason to take the model seriously.

Whereas Experiments 1-3 examined the ability of our model to account for the speed and accuracy of behavioral change detection judgments for scene pairs, Experiment 4 provided a further test by asking if our model could account for patterns of brain activity recorded while a single scene was being held in working memory. This is important because a variety of decision processes are involved in making same-different judgments of scene pairs, and we wanted to ensure that our model could account for the working memory representations themselves and not just for the same-different decision processes. To accomplish this, we recorded the electroencephalogram (EEG) while participants performed a working memory task with real-world scenes, and we used representational similarity analysis (Kriegeskorte et al., 2008) to determine whether differences in the pattern of neural activity during the delay period of the task for different scenes could be accounted for by differences in the vectors produced by those scenes in the model.

An additional goal of the present study was to assess the ability of the different areas within CORnet to predict behavioral performance and account for the neural data. That is, by assessing the ability of each individual area of CORnet to predict behavior and neural activity, we were able to test the contributions of spatially detailed, V1-like representations and more abstract, IT-like representations to working memory. For the sake of simplicity, we focus primarily on the V1_COR_ and IT_COR_ areas of CORnet in the main text, with V2_COR_ and V4_COR_ results provided in supplementary materials. However, when comparing our theory with the lower reference point (RGB vector distances) and upper reference point (perceptual similarity ratings), we focus on the amount of variance explained by the full model, including all four areas of CORnet. This is because the RGB bitmaps contain all of the information needed to perform the task while the similarity ratings potentially include information from the entire brain, so it would be inappropriate to compare the amount of variance accounted for by the bitmaps and by the similarity ratings with the amount of variance accounted for by only two of the four areas of our model.

## Transparency and Openness

In the sections below, we report how our sample size was determined along with all data exclusions, experimental protocols, and measures. The experiments were not preregistered, but we provide information below about post hoc decisions that were made in the analysis process. All data, analysis code, stimulus presentation code, and research materials are freely available at https://osf.io/7ukgj/. This will make it possible for other researchers to compare alternative models against our model using the substantial sets of data we collected (once alternative models of working memory for natural scenes are developed).

## Experiment 1: Predicting Behavioral Change Detection Performance with a Flicker Task

The aim of Experiment 1 was to assess the population vector model’s ability to account for variability in behavioral change detection performance with the flicker change detection task depicted in Figure 3a and the scenes illustrated in Figure 3c. A unique pair of scenes was chosen on each trial, and observers saw an alternating series of these two scenes. The two scenes were usually different, and observers simply pressed a button when they detected the difference. The amount of time required to detect the difference should be greater for more similar scenes, but the effective similarity of a given pair of scenes depends on how those scenes are represented in the brain. The population vector model represents each scene as a vector within CORnet, so the similarity between the scenes—and hence the response time—should be predicted by the distance between the vectors for the two scenes. That is, we predicted that the response time (RT) would increase as the vector distance decreased.

**Figure 3.**
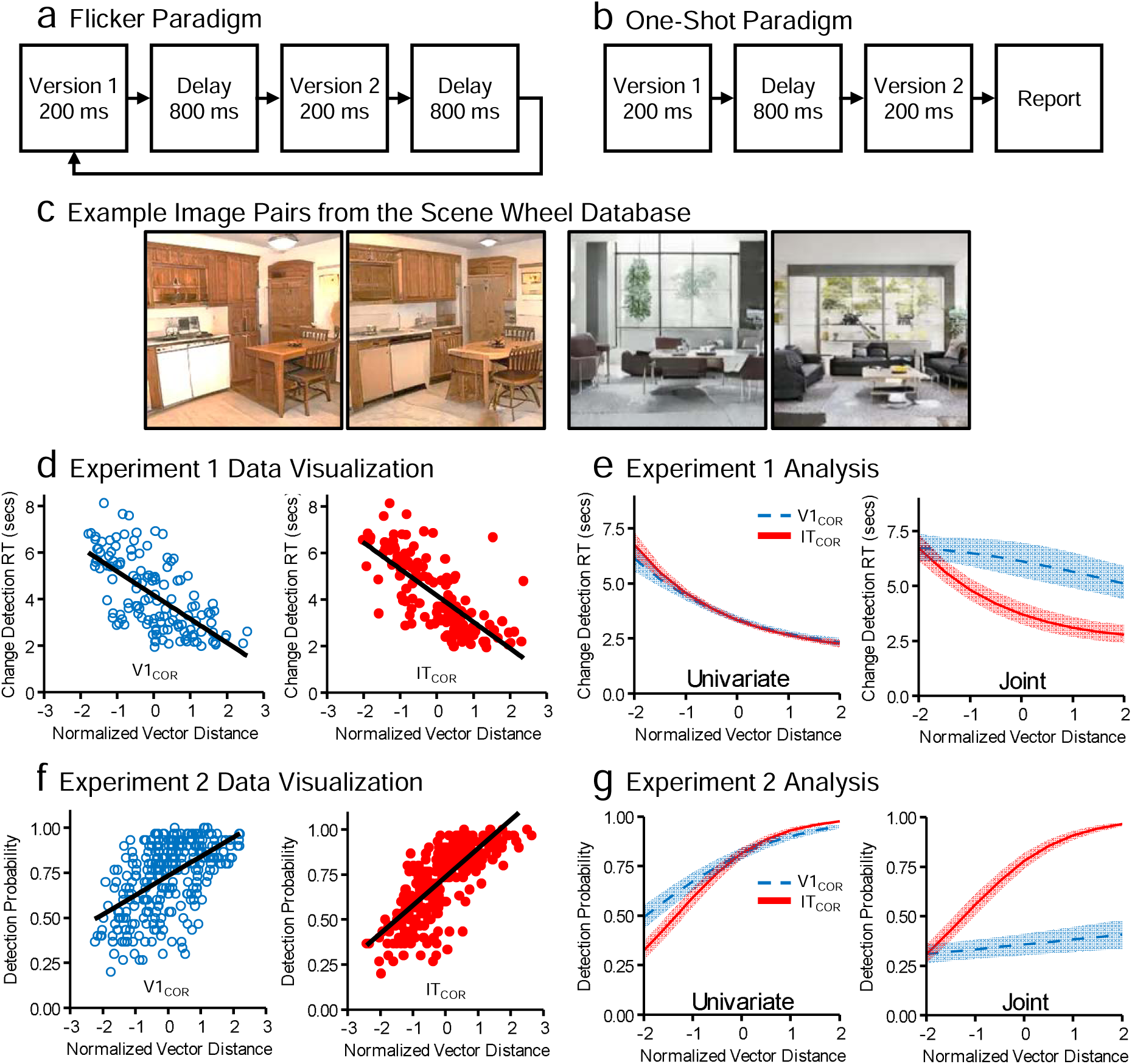
Tasks and Primary Results from Experiments 1 and 2. *Note.* **a**, Flicker procedure used in Experiment 1. Two versions of a scene alternate until the participant detects a difference. **b**, One-shot procedure used in Experiment 2. Each of the two scene versions is presented once, and the participant then reports whether they were identical or different. **c**, Examples of two pairs of images from the scene database. The images in the left pair are subtly different from each other, whereas the images in the right pair are quite different. **d**, Scatterplot visualization of the results from Experiment 1, showing the relationship between either the V1_COR_ or IT_COR_ vector distances for a given scene pair and the mean response time (RT) across participants for that pair. Changes were correctly detected on 94.52% of change trials, with a mean RT of 4050 ms. Changes were reported on 11.80% of no-change trials (not shown), with a mean RT of 6578 ms. **f**, Scatterplot visualization of the results from Experiment 2, showing the relationship between either the V1_COR_ or IT_COR_ vector distances for a given scene pair and the proportion of participants who detected a change for that pair. A change was reported on 19.97% of no-change trials (not shown). **e & g**, Results of the statistical analysis of the single-trial RTs from Experiment 1 (**f**) and the single-trial binary responses from Experiment 2 (**g**). The univariate plots show predicted values when V1_COR_ and IT_COR_ were analyzed separately, whereas the joint plots shows predicted values when both variables were included simultaneously (with each effect being centered at -2 SD on the other variable). Shading indicates +/-1 SE.

The scenes in this experiment were taken from a database of computer-generated naturalistic images of real-world scenes that differ systematically from each other in a continuous high-dimensional scene space (Son et al., 2022). As illustrated in Figure 4, the scenes were selected from a circular similarity space, which avoids guessing artifacts that can occur with bounded stimulus spaces (Thiele et al., 2011). We randomly selected pairs of scenes from a set that ranged from identical to modestly different in terms of their distance in the circular similarity space (see examples in Figure 3c). A given pair of scenes differed in features that were spread broadly across the images; this property was well suited for CORnet, which has equal acuity across space rather than containing a central high-acuity region (a fovea). Experiment 3 will examine scene pairs that differ in the presence or absence of a single object.

**Figure 4.**
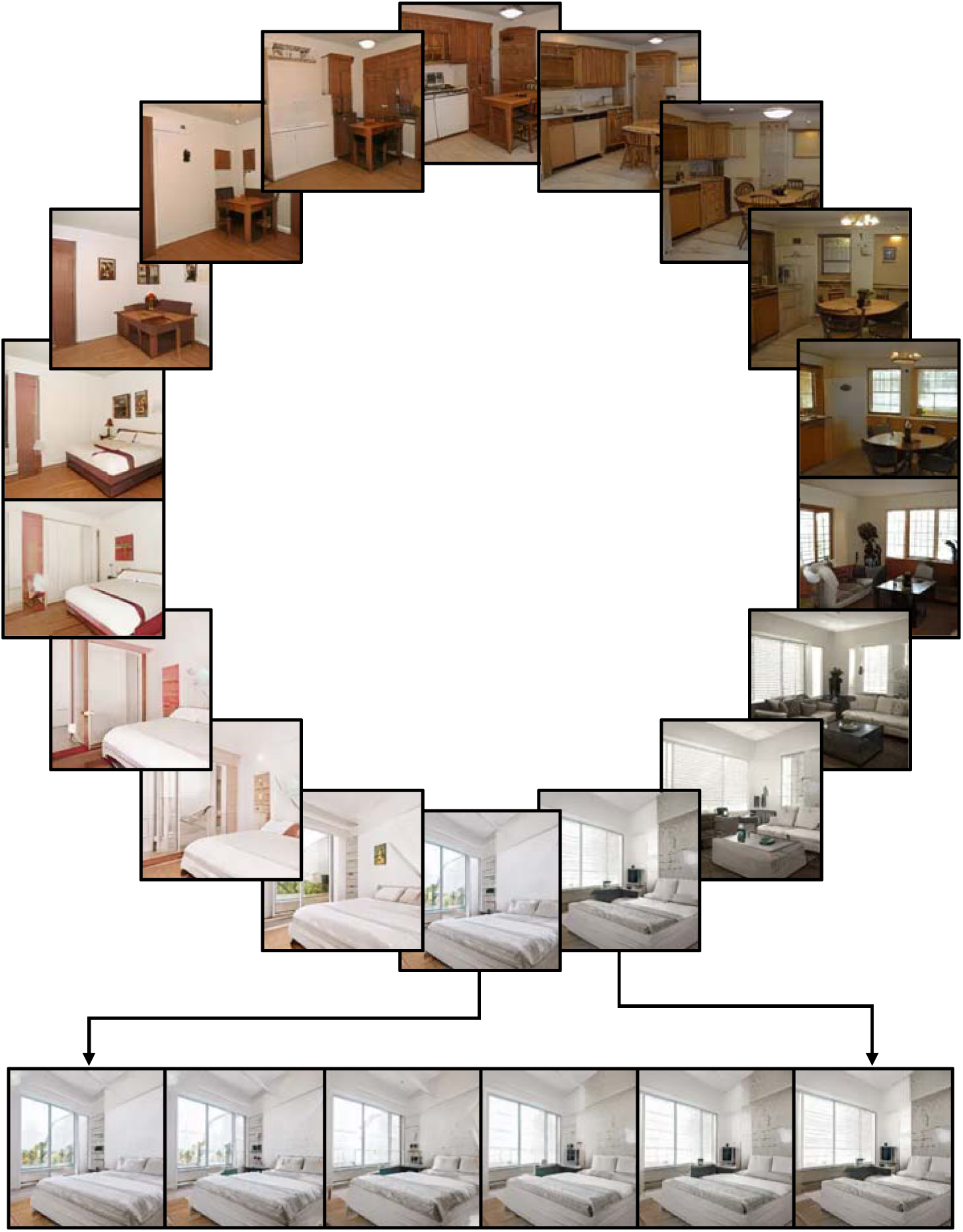
Example of the scene wheels used in Experiments 1 and 2. *Note*. This circle of images shows one image for every 18° of rotation around the multidimensional space for one of the scene wheels. The line of images along the bottom shows one image for every 3° of rotation. The actual stimulus set contains one image for every 1° of rotation. In Experiment 1, a given pair of images was separated by 4, 8, or 16° (with a random starting point). In Experiment 2, a given pair of images was separated by 6, 12, or 18° (with a random starting point). The image generation process is described in reference 47, and the images were obtained from https://osf.io/h5wpk/.

Human-rated measures of similarity were obtained from a separate group of observers who were shown each scene pair side by side and asked to make untimed ratings of their visual similarity. As described earlier, the rated similarity between a pair of scenes should strongly predict the RT for determining that the two images are different in the change detection task, serving as an upper reference point for the model’s ability to predict change detection RTs.

## Method

### Participants

All participants in all experiments presented here reported having normal or corrected-to-normal visual acuity, normal color vision, and a lack of neurological problems. All experiments reported here were also approved by the UC-Davis Institutional Review Board, and all participants provided informed consent before data collection began. For Experiments 1 through 3, power analyses based on pilot data indicated that 80% power could be achieved with fewer than 10 participants for the main analyses. Nonetheless, we aimed for an *N* of 30 to allow more precise parameter estimates and more sophisticated statistical modeling. Consequently, 30 participants were recruited for Experiment 1. An additional group of nine participants was recruited from the same population to perform a similarity rating task.

The participants were UC-Davis college students who received course credit. Table 1 provides the demographic information for all four experiments. Race and ethnicity demographics were defined using the approach required by the National Institutes of Health when the data were collected (2022-2024), which involves one multiple-choice question about race (American Indian or Alaska Native, Asian, Black or African American, Native Hawaiian or Other Pacific Islander, White) and another about ethnicity (Hispanic or Latino, Not Hispanic or Latino). Gender information was collected using a three-option multiple-choice question (Man, Woman, Non-Binary\Third Gender). Participants were also given the option of reporting more than one race and the option of declining to report race, ethnicity, and/or gender.

**Table 1.** Demographics for each experiment.

| Experiment # | Age Range | Gender Breakdown | NIH Race\Ethnicity Categories |
| --- | --- | --- | --- |
| 1<br>(Main Experiment) | 18 – 29<br>(Mean = 20.2, SD = 2.08) | 19 Women, 10 Men<br>1 Did not Report (DNR) | 16 Asian, 1 Black, 2 Native American, 2 Did Not Report, 2 Hispanic, 7 White |
| 1<br>(Similarity Ratings) | 20 – 22<br>(Mean = 20.39, SD = 0.62) | 4 Women, 4 Men,<br>1 Nonbinary | 7 Asian, 2 White |
| 2 | 18 – 23<br>(Mean = 19.4, SD = 1.18) | 26 Women, 3 Men<br>1 Did Not Report | 14 Asian, 1 Black, 4 Hispanic, 3 Did Not Report, 8 White |
| 3 | 18 – 21<br>(Mean = 19.7, SD = 0.95) | 28 Women, 6 Men | 18 Asian, 1 Black, 1 Native American, 3 Hispanic, 2 Did Not Report, 9 White |
| 3<br>(Similarity Ratings) | 18 - 21<br>(Mean = 19.60, SD = 1.12) | 7 Women, 3 Men | 5 Asian, 1 Hispanic, 2 White, and 1 Did Not Report |
| 4 | 18 – 29<br>(Mean = 21.03, SD = 2.12) | 23 Women, 4 Men, 2<br>Nonbinary | 16 Asian, 1 Black, 1 Native American, 4 Hispanic, 4 White, 3 Did Not Report |

### Stimuli

The stimuli were presented using PsychoPy (Peirce, 2009) on an HP ZR2440W LCD monitor with a gray background at a viewing distance of 100 cm in a controlled laboratory environment. The images were selected from a database of images created by generative adversarial networks (GANs) and designed to vary systematically in a high-dimensional space (Son et al., 2022). Each image was a photorealistic depiction of a typical indoor scene (see examples in Figures 3c and 4). From the high-dimensional space, scenes were selected that varied gradually and systematically around a 2-dimensional circular wheel within the high-dimensional space (a *scene wheel*). The database contained five such wheels, each with 360 scenes at each of 5 radii. We used images from the largest radius of each of the five wheels (see example wheel in Figure 4). Each image subtended 16° × 16° of visual angle.

### Procedure

A total of 165 image pairs from these scene wheels were used with the flicker task shown in Figure 3a. For each pair, one image was first selected randomly from among all the images contained in outer radii of the five scene wheels. For 80% of pairs, the other image for the pair was ±4, ±8, or ±16 degrees away (with equal likelihood) from the first image on the same wheel. On the remaining 20% of pairs, the second image was identical to the first image. Each participant received one trial with each of the 165 image pairs in random order.

On each trial, the two images from a pair alternated, with a duration of 200 ms per presentation and a blank gap of 800 ms between presentations. Participants were instructed to make a speeded response on a keyboard with the index finger of the preferred hand as soon as they detected a difference between the two scenes (or to withhold responding if they detected no difference). The sequence terminated when the participant responded or after seven cycles of the two scenes, whichever came first. Before beginning the experiment, each participant completed a practice session (with images not used in the main task) that continued until they correctly classified 10 out of 10 change-vs-no-change image pairs in a row. The stimuli and experimental control script are available at https://osf.io/7ukgj/.

### Computation of DNN Vectors for each Scene

In all four experiments, we used the “skip” variant of CORnet (*CORnet-S*) (Kubilius et al., 2018) as our model of the perceptual representation of a scene to compute the population activation vector for each scene. The architecture of CORnet-S combines elements of the original AlexNet architecture (Krizhevsky et al., 2012) and ResNet (K. He et al., 2016). The input is a 224×244 pixel image, with red, green, and blue values at each pixel. The model is divided into four areas (V1_COR_, V2_COR_, V4_COR_, and IT_COR_), each consisting of a sequence of multiple layers that simulate recurrent processing via unrolling into a sequence of feedforward layers (Liao & Poggio, 2020). Each subsequent layer within an area is created via a sequence of convolution, normalization, and rectified linear unit (ReLU) operations. The output of an area passes through an additional maxpooling nonlinearity before becoming the input to the next area. The weights were set via a supervised learning procedure in which the model was trained to classify a massive set of labeled photographs of real-world objects (Russakovsky et al., 2015). During training, the IT_COR_ output layer was fed into a linear classifier, which used the pattern of activity to categorize the identity of the image.

We downsampled the photographic images used in the present study to 224 × 224 and fed them into the pretrained version of CORnet-S (https://github.com/dicarlolab/CORnet). For each scene, we used the vector of ReLU values (which are analogous to firing rates) of the final layer of a given area to model the perceptual representation in that area. We assume that the working memory representation of a scene is a weak (scaled-down) version of this pattern. However, scaling the values does not change the pattern of scene-to-scene similarity, so we did not actually scale the values in our analyses. We simply computed the Euclidean distance between the vectors for each pair of scenes and used that distance to predict behavioral change-detection performance for that pair of scenes.

As an a priori modeling decision, we applied a normalization operation to the vector distances that was inspired by Weber’s law, which states that the perceived difference between two values is proportional to the magnitude of the values (Fechner, 1948). For example, the ability to perceive a difference in weight between two objects weighing 100 and 110 g is approximately the same as the ability to perceive a difference in weight between two objects weighing 10 and 11 g. One potential explanation for this law is that the variance in neural firing rates increases proportionately with the mean firing rate (Tolhurst et al., 1981), often with a lognormal distribution (Nieder & Miller, 2003; Roxin et al., 2011) (although many other explanations have been proposed; Deco & Rolls, 2006; Pardo-Vazquez et al., 2019; Simen et al., 2016). Weber’s law also appears to apply to working memory performance, at least in the case of detecting changes in location (Palmer, 1986). We implemented the normalization procedure inspired by Weber’s law in our model by dividing the distance between the vectors for a pair of scenes by the vector length for the first-presented scene.

Note that this normalization operation is somewhat similar to the divisive normalization that produces capacity limitations in some models of visual working memory (Bays, 2014; Bouchacourt & Buschman, 2019; Wei et al., 2012), and it may have a similar effect: A scene that strongly activates more units will have a longer vector length, and a larger absolute change will be needed to reliably detect a difference between scenes with longer compared to shorter vector lengths. However, the purpose of the normalization operation in the present model was to implement the essential insight of Weber’s law rather than to account for capacity limitations. Indeed, the concept of capacity limitations is not readily extendable to natural scenes that do not have an unambiguous set of simple, discrete objects. Note also that, as described in the section entitled *Effect of Normalization across all three Experiments*, our normalization approach had very little impact on the vector distances for the image pairs used here. Thus, although our normalization approach might have a meaningful impact in future experiments, it is of minimal importance in the present study and is included because it was an a priori decision.

### Computation of RGB Vectors for each Scene

The RGB vectors were obtained using an approach very similar to that used to compute the DNN vectors. Each image in this study was an X × Y × RGB bitmap with X and Y indicating the horizontal and vertical dimensions, respectively, and RGB being the triplet of intensity values for the three color channels (Red, Green, Blue) at a given location. This is analogous to the array of activation values for a given layer of CORnet, and the vector distance for a given pair of images was computed just as we did with the CORnet vectors, as the Euclidean distance between the vectors for each pair of scenes. For the sake of consistency, we then applied our Weber-motivated normalization operation to these distances.

### Analytic Approach

Trials were excluded from the statistical analysis if the two stimuli were identical (i.e. catch trials), if the participant failed to respond, or if the RT was <200 ms (relative to the onset of the second image in the trial, because changes could not be detected prior to the second image). The single-trial, single-participant RTs from the included trials were analyzed using a generalized linear mixed effects (GLME) analysis with PROC GLIMMIX in SAS 9.3 using a lognormal identity link to account for the non-Gaussian distribution of RT residuals. Quadratic terms were included for all predictors to account for the fact that RTs are bounded at the lower end^2^. Random intercepts for subjects and items were specified. Random slopes were tested but not included because they either did not significantly improve the fit of the statistical model or resulted in convergence failures when included. In the initial analyses, the RTs from each trial predicted by the normalized vector distance between the two images used on that trial, with separate analyses for V1_COR_ and IT_COR_. These initial analyses allowed us to ask how well the vector distances in a single area could predict the observed RTs.

We followed the initial analyses with follow-up analyses designed to ask how well the activation vectors in a given area predicted RT when another variable was controlled. For example, we entered both the V1_COR_ and IT_COR_ distances into the same statistical analysis so that we could ask how well the V1_COR_ distances predicted RT when the IT_COR_ distances were controlled and how well the IT_COR_ distances predicted RT when the V1_COR_ distances were controlled. The amount of variance uniquely explained by one of the two distance variables was quantified via pseudo-*r^2^* values, which were computed by estimating the decrease in unexplained item-level variance when the predictor of interest was added to the analysis. We used the same approach to ask whether the distances in either V1_COR_ or IT_COR_ remained predictive of RT when ratings of perceptual similarity were controlled.

As explained earlier, most of the analyses in the main text focused on V1_COR_ and IT_COR_ distances. We also performed analyses including all four areas (the *full model*) so that we could assess the predictive value of the entire population vector model (the overall r^2^) and compare this with the r^2^ for the upper and lower reference points. However, we refrained from attempting to assess the unique variance accounted for by each of the four areas within this analysis because of the substantial collinearities present in the full model (which included both linear and quadratic terms for all four areas), which almost certainly resulted in complicated covariance patterns that would potentially lead to misleading estimated effects for the individual areas.

### Human Similarity Ratings

For each image pairs used in the main experiment, we obtained similarity ratings from a separate group of participants. On each trial of this task, one pair of images was selected at random from the full set, and both images were shown simultaneously side by side. One image (selected at random from the pair) was centered 5° to the left of fixation, with the other being centered 5° to the right of fixation. Each image subtended 9° × 9° of visual angle. We also included the pairs from no-change trials, which were identical. Each participant was asked to rate the degree of difference between the two images on a scale of 0 to 100 (from maximally similar to maximally dissimilar) using an on-screen sliding scale. This repeated until all pairs had been rated. In this manner, each of the nine raters produced a rating for every image pair presented in the main experiment. Participants were given unlimited time to make each rating, and the pair of images remained visible until the rating was given.

Because some experience was necessary for participants to calibrate their perceptions of similarity to the rating scale, we had participants rate all the images twice in two completely distinct passes (i.e., a practice pass followed by the actual rating task). Ratings from the second pass were then averaged across participants for each image pair, with the produced mean scores being used as our measure of human-judged similarity. To assess the consistency of this measure, we computed the intraclass correlation (ICC(C,k)) of the ratings (McGraw & Wong, 1996).

## Results and Discussion

### Overall behavioral performance

Changes were correctly detected on 94.45% of change trials, with a mean RT of 4052 ms. Changes were reported on 11.17% of no-change trials (not analyzed here), with a mean RT of 6578 ms.

### Ability of the model to predict change detection performance

Figure 3d shows scatterplot visualizations of the results from the main experiment for areas V1_COR_ and IT_COR_ (see Supplementary Figure S1 for the V2_COR_ and V4_COR_ results). Each point represents the mean RT across participants for a given scene pair. As predicted, RTs became progressively faster as the vector distance increased. This pattern was present for both V1_COR_ and IT_COR_, although somewhat stronger for IT_COR_. Note that the scatterplots in Figure 3d are just visualizations, and our GLME statistical analyses examined the single-participant, single-trial RTs rather than the mean RTs across participants.

The initial statistical analyses separately tested whether the V1_COR_ or IT_COR_ vector distances for a given scene pair could predict the single-trial RTs for that pair. As illustrated in Figure 3e, significant effects were observed for both V1_COR_ (r^2^ = .44) and IT_COR_ (r^2^ = .57). The test statistics corresponding to the r^2^ values for single predictor variables are shown in Table 2, with all data and analysis code provided at https://osf.io/7ukgj/.

**Table 2.**
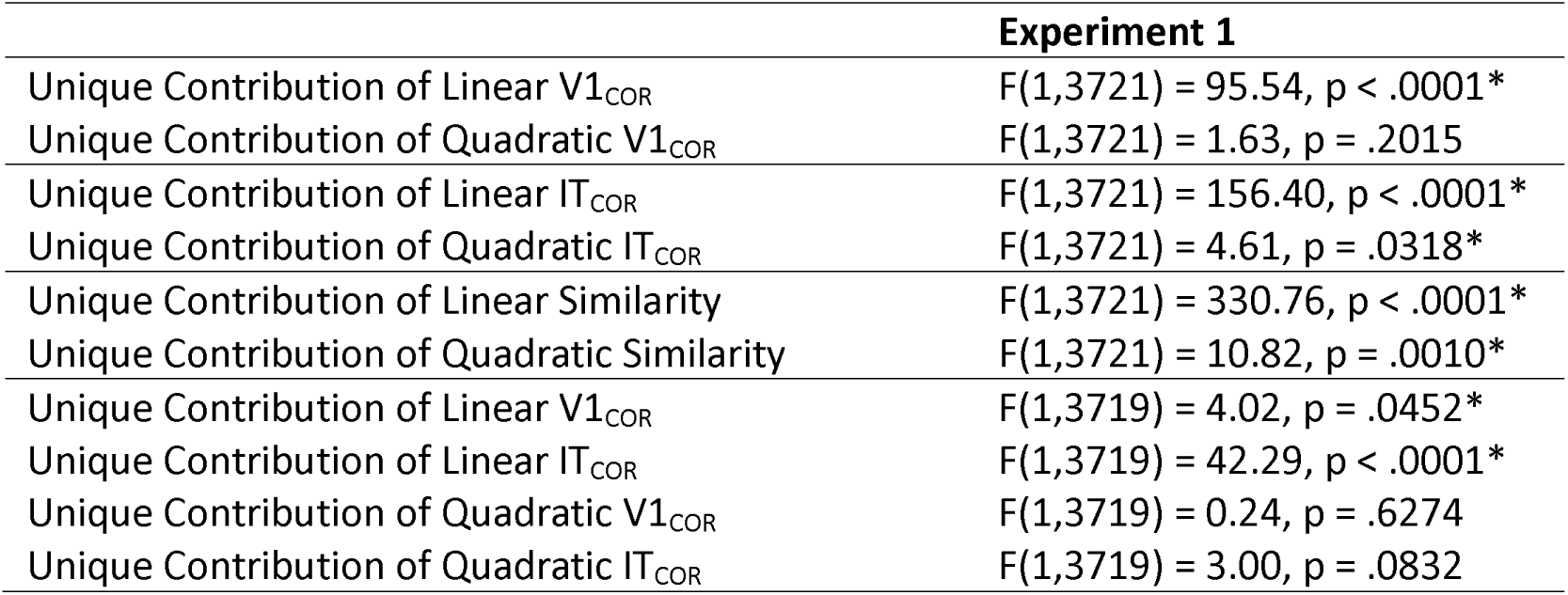
Test statistics for analyses of single-trial response times in Experiment 1.

|  | **Experiment 1** |
| --- | --- |
| Unique Contribution of Linear $V1_{COR}$ | $F(1,3721) = 95.54, p < .0001^*$ |
| Unique Contribution of Quadratic $V1_{COR}$ | $F(1,3721) = 1.63, p = .2015$ |
| Unique Contribution of Linear $IT_{COR}$ | $F(1,3721) = 156.40, p < .0001^*$ |
| Unique Contribution of Quadratic $IT_{COR}$ | $F(1,3721) = 4.61, p = .0318^*$ |
| Unique Contribution of Linear Similarity | $F(1,3721) = 330.76, p < .0001^*$ |
| Unique Contribution of Quadratic Similarity | $F(1,3721) = 10.82, p = .0010^*$ |
| Unique Contribution of Linear $V1_{COR}$ | $F(1,3719) = 4.02, p = .0452^*$ |
| Unique Contribution of Linear $IT_{COR}$ | $F(1,3719) = 42.29, p < .0001^*$ |
| Unique Contribution of Quadratic $V1_{COR}$ | $F(1,3719) = 0.24, p = .6274$ |
| Unique Contribution of Quadratic $IT_{COR}$ | $F(1,3719) = 3.00, p = .0832$ |

Unsurprisingly, the V1_COR_ and IT_COR_ vector distances were correlated (r = .745, p < .001). We therefore performed a GLME analysis that included both the V1_COR_ and IT_COR_ vector distances as joint predictors, allowing us to estimate the unique contribution of each area (Figure 3e, right). Both V1_COR_ and IT_COR_ explained unique variance in the change detection RTs with the effect being numerically larger for IT_COR_ (r^2^ = .15) than for V1_COR_ (r^2^ = .02).

To assess the full predictive ability of the population vector model, we asked how well a GLME analysis including the vector distances from all four areas of CORnet could predict RT. This full model was strongly predictive (r^2^ = .76, χ^2^(8) = 168.93, p < .0001). The fact that the model accounted for 76% of the item-level variance across scene pairs is remarkable for single-trial RT data and provides initial evidence that our model is viable.

### Ability of RGB bitmaps to predict change detection performance

Our next step was to ask how well change detection RTs could be explained by vector distances obtained directly from the RGB bitmaps for the image pairs. This served as a lower reference point: Any useful model should be able to explain behavioral performance substantially better than the RGB bitmaps. We found that, on their own, the vector distances from the RGB bitmaps could account for 29% of the variance across image pairs (r^2^ = .29, χ^2^(2) = 43.07, p < .0001). This is less than the variance accounted for individually by V1_COR_ (r^2^ = .44) and by IT_COR_ (r^2^ = .57), and it is much less than the variance accounted for by the full model containing all four areas of CORnet (r^2^ = .76). Thus, our model readily passed this minimal test.

### Ability of similarity ratings to predict change detection performance

We next asked how well the perceptual similarity ratings for each scene pair predicted the change detection RTs. The intraclass correlation indicated that the similarity ratings were highly consistent across observers (r(164, 1312) = .948, p < .0001). When entered into a GLME analysis as the sole predictor of RT, these ratings explained 78% of the variance in RT across scene pairs (χ^2^(2) = 178.03, p < .0001). Note that the combination of a very high intraclass correlation and very strong prediction of change detection performance indicates that our procedure of averaging ratings across 9 observers produced a robust index of perceptual similarity for the scene pairs.

We assumed from the outset that the perceptual similarity ratings should provide an excellent prediction of behavior, serving as a reasonable upper reference point for the amount of variance explained by our model. The fact that the variance accounted for by the full model (76%) was nearly as large as the variance accounted for by the perceptual similarity ratings (78%) indicates that our model explains behavioral performance about as well as could be reasonably expected. It is possible that a different method for obtaining perceptual similarity ratings would allow these ratings to explain more than 78% of the RT variance. However, even if improved similarity ratings accounted for 99% of the item-level RT variance, the fact that the model accounted for 76% of the item-level RT variance would still indicate that the model is performing very well. Moreover, the fact that distance vectors from the RGB bitmaps accounted for only 29% of the item-level variance indicates that it is not trivial for a representation of the scenes to account for a large percentage of the variance.

### Unique predictive ability of the model versus the similarity ratings

We also asked whether our model could account for variance in behavior that was not captured by the similarity ratings. This was a high bar given that—unsurprisingly— similarity ratings were negatively correlated with V1_COR_ and IT_COR_ vector distances (r(132) = -.61, p < .001 and r(132) = -.75, p < .001, respectively), such that higher ratings of similarity were associated with smaller vector distances. This collinearity impacts the interpretability of individual predictors, so we focused instead on the predictive power of the full model including all four layers of CORnet. A statistical analysis including these four areas along with the similarity ratings strongly predicted RT (r^2^ = .84, χ^2^(10) = 214.07, p < .0001). After controlling for the similarity ratings, the four CORnet areas together still accounted for a significant proportion of the RT variance (r^2^ = .07, χ^2^(8) = 36.04, p < .0001). Likewise, the similarity ratings accounted for a significant amount of variance in RT after controlling for the full CORnet model (r^2^ = .09, χ^2^(2) = 45.14, p < .0001).

Because some of our previous analyses included only V1_COR_ and IT_COR_, we conducted an additional set of analyses using just these two areas along with the similarity ratings. This statistical analysis accounted for significant item-level RT variance overall (r^2^ = .81, χ^2^(6) = 191.87, p < .0001). After controlling for the similarity ratings, the two CORnet areas together accounted for a small but significant proportion of RT variance after controlling for similarity (r^2^ = .03, χ^2^(4) = 13.84). In this analysis, the similarity ratings accounted for a moderate amount of RT variance after accounting for the two areas of the model (r^2^ = .21, χ^2^(2) = 82.51, p < .0001). The unique variance explained by V1_COR_ and IT_COR_ in isolation after controlling for similarity ratings is provided in supplementary Table S1.

The key conclusion of these analyses is that our model accounts for a small but significant amount of variability in change detection RT even after controlling for the similarity ratings. One might have expected that exactly the same representations would be used for change detection and for similarity rating, in which case the model should have accounted for no unique variance. Indeed, the model would have still been valuable if it had explained no unique variance. The fact that it does explain some unique variance provides additional reason to take it seriously as a model of working memory. The General Discussion will consider the question of how the model could explain unique variance.

## Experiment 2: Extension to a One-Shot Change Detection Paradigm

Most change detection studies testing quantitative models of visual working memory have used a *one-shot* paradigm, in which a stimulus is shown only once on each trial, and accuracy rather than RT is the primary dependent variable. In Experiment 2, we used this approach with the same stimulus database used in Experiment 1 (but with more dissimilar scene pairs to prevent floor effects). As shown in Figure 3b, observers saw one scene from a pair followed by a delay and then the other scene, and then they made an unspeeded change/no-change response. We predicted that the probability of detecting a change for a given scene pair would be proportional to the vector distance between the two scenes.

### Method

Thirty new participants were recruited from the same population for this experiment (see Table 1 for demographic information). Experiment 2 was otherwise identical to Experiment 1 except as noted here.

The stimuli were drawn from the same database used in Experiment 1, but with image pairs separated by ±6, ±12, or ±18 degrees on change trials (because the one-shot task was more difficult than the flicker task). The sample and test images were identical on 25% of trials. On each trial, one image from a pair was presented for 200 ms, followed by a blank screen for 800 ms and then the other image for 200 ms. Participants were instructed to make an unspeeded keyboard response after the test image to indicate whether the two images were identical (index finger of the preferred hand) or were different (index finger of the non-preferred hand).

Because each trial was brief and the dependent variable was binary (correct or incorrect) rather than continuous (RT), we increased the number of trials (and therefore the number of scene pairs) to 400. We analyzed the single-trial, single-subject binary responses from the change trials (coded as 1 for correct and 0 for incorrect) using the same GLME analysis approach used in Experiment 1, except specifying a binary distribution and logit link.

## Results and Discussion

### Overall behavioral performance

A change was reported correctly on 73.4% of change trials. A change was reported incorrectly (i.e., a false alarm) on 22.1% of no-change trials (not analyzed further).

### Ability of the model to predict change detection performance

Figure 3f shows scatterplot visualizations of the results for V1_COR_ and IT_COR_ (see Supplementary Figure S1 for the V2_COR_ and V4_COR_ results). Each point reflects the proportion of participants who detected the change for a given scene pair. As predicted, the probability of detecting a difference was greater for pairs with larger vector distances, especially for IT_COR_.

The single-participant, single-trial binary responses on change trials were initially analyzed in separate GLME analyses for V1_COR_ and IT_COR_. As illustrated in Figure 3g, change detection was strongly and significantly predicted by the IT_COR_ vector distances (r^2^ = .71) and approximately half as strongly by the V1_COR_ vector distances (r^2^ = .36). The test statistics corresponding to the r^2^ values for individual predictor variables are shown in Table 3.

**Table 3.**
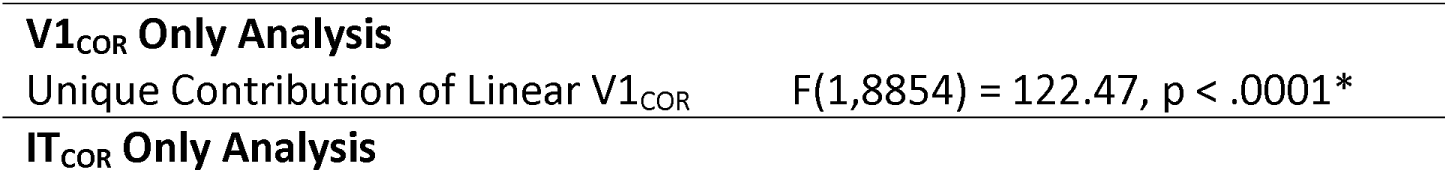
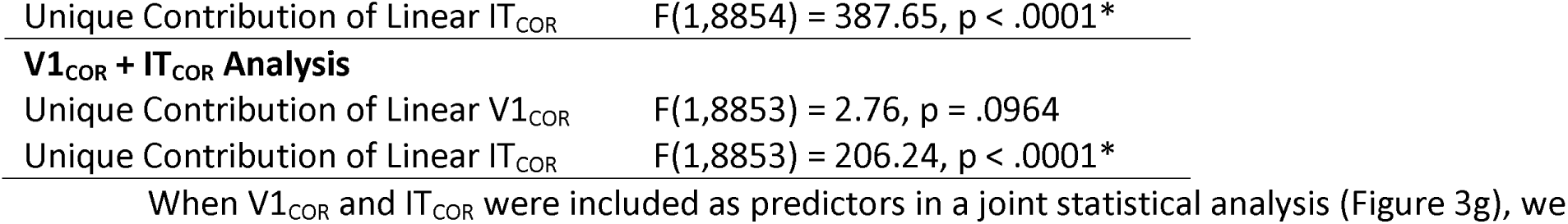
Test statistics for analyses of single-trial accuracy (Experiment 2)

|  |  |
| --- | --- |
| **$V1_{COR}$ Only Analysis** |  |
| Unique Contribution of Linear $V1_{COR}$ | $F(1,8854) = 122.47, p < .0001^*$ |
| **$IT_{COR}$ Only Analysis** |  |
| Unique Contribution of Linear $IT_{COR}$ | $F(1,8854) = 387.65, p < .0001^*$ |
| **$V1_{COR}$ + $IT_{COR}$ Analysis** |  |
| Unique Contribution of Linear $V1_{COR}$ | $F(1,8853) = 2.76, p = .0964$ |
| Unique Contribution of Linear $IT_{COR}$ | $F(1,8853) = 206.24, p < .0001^*$ |

When V1_COR_ and IT_COR_ were included as predictors in a joint statistical analysis (Figure 3g), we found that these two variables together accounted for 72% of the item-level variance in the probability of change detection (χ^2^(2) = 277.06, p < .0001). Within this analysis, IT_COR_ explained significant unique variance (r^2^ = .35), but V1_COR_ did not (r^2^ < .01). A full model including vector distances from all four CORnet areas explained 80% of the item-level variance in the single-trial binary responses (χ^2^(8) = 336.99, p < .0001).

### Ability of RGB bitmaps to predict change detection performance

When we computed vector distances from the RGB bitmaps to provide a lower reference point for variance explained, we found that these vector distances accounted for 29% of the item-level variance (χ^2^(1) = 84.67, p < .0001). Our model of working memory clearly outperformed this minimal level of predictive ability. Because this experiment used 400 different image pairs (as opposed to 165 pairs in Experiment 1), it was not feasible to obtain perceptual similarity ratings for this task to serve as an upper reference point. However, even if similarity ratings explained 99% of the item-level variance, the fact that the full version of our model of working memory explained 80% of the item-level variance (or 72% if only V1_COR_ and IT_COR_ were included) indicates that this model is viable.

### Experiment 3: Replication with an Object Addition-Deletion Change Detection Paradigm

The scene pairs used in Experiments 1 and 2 differed broadly across the images, making them particularly well suited for CORnet, which has no fovea and instead has a uniform input resolution across spatial locations. However, most prior research using change detection with natural scenes has examined focal changes, such as the addition or deletion of an object from a scene (Hollingworth & Henderson, 2002; Rensink et al., 1997). Experiment 3 therefore examined the ability of our model to predict performance with scene pairs that differed solely in the presence or absence of a single object, as illustrated in Figure 5a. The flicker paradigm used in Experiment 1 was also used in this experiment.

**Figure 5.**
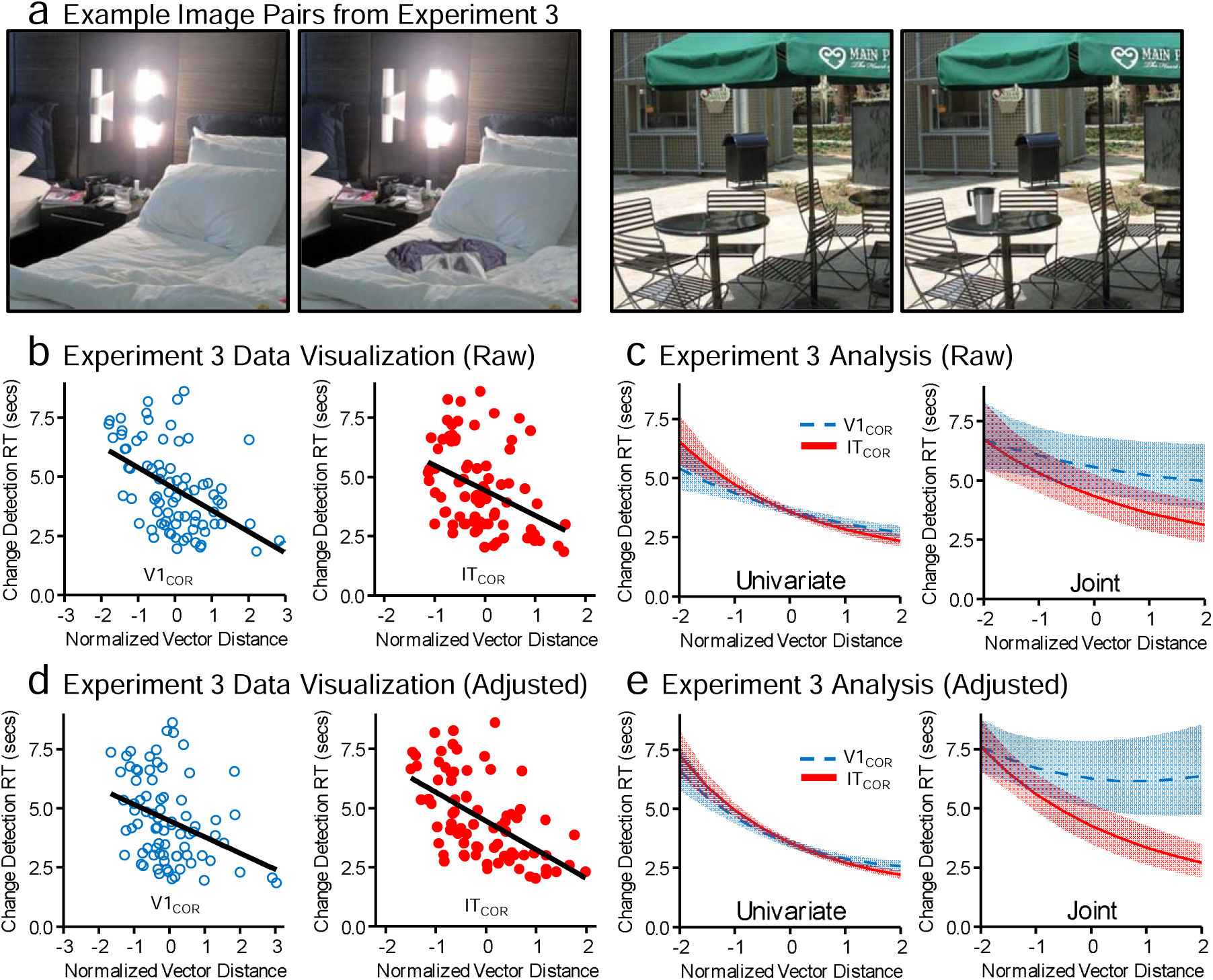
Tasks and Primary Results from Experiments 3. *Note.* **a**, Examples of two pairs of images used in Experiment 3. The images in a pair differ solely in the presence of absence of a single object. **b**, Scatterplot visualization of the results from Experiment 3, showing the relationship between either the V1_COR_ or IT_COR_ vector distances for a given scene pair and the mean response time (RT) across participants for that pair. Changes were correctly detected on 90.81% of change trials, with a mean RT of 4289 ms. Changes were reported on 16.48% of no-change trials (not shown), with a mean RT of 6473 ms. **c**, Results of the statistical analysis of the single-trial RTs from Experiment 3. Shading represents the standard error. The univariate plots show predicted values when V1_COR_ and IT_COR_ were analyzed separately, whereas the joint plot shows predicted values when both variables were included simultaneously (with each effect being centered at -2 SD on the other variable). **d, e**, Same as **b** and **c**, but using vector distances computed after adjusting for the distance from the center of the image. Shading indicates +/- 1 SE.

## Method

Thirty-four new participants were recruited for this experiment (see Table 1 for demographic information). Experiment 3 was identical to Experiment 1 except as noted here.

For this experiment, we created a custom set of image pairs that differed in the presence or absence of a single object. The base images were selected from the COCO database (Lin et al., 2015). An object was then added to a copy of each scene using Photoshop. The objects were selected from the BOSS database (Brodeur et al., 2014), were chosen to match the content of the scene and were sized and located to look natural within it. Detailed information about the sizes and locations of the objects are provided in Supplementary Table S2. The resulting images and the corresponding item masks are available at https://osf.io/7ukgj/. On each trial, one pair of scenes was randomly selected and presented using the flicker paradigm as in Experiment 1. The original version or the Photoshopped version was equally likely to appear first. The sample and test images were identical on 10% of trials. Each participant received a total of 88 trials in random order, with different randomly selected scene pairs on each trial.

As in Experiment 1, we obtained perceptual similarity ratings for each image pair in an additional sample of 11 participants (see Table 1 for demographic details).

## Results and Discussion

### Overall behavioral performance

Changes were correctly detected on 90.77% of change trials, with a mean RT of 4289 ms. Changes were reported on 16.23% of no-change trials (not analyzed here), with a mean RT of 6473 ms.

### Ability of the model to predict change detection performance

Figure 5b shows scatterplot visualizations showing the relationship between the mean RT for each scene (averaged over participants) and the corresponding vector distances in V1_COR_ and IT_COR_ (see Supplementary Figure S1 and Supplementary Table S3 for the V2_COR_ and V4_COR_ results). As in Experiment 1, RTs were faster for pairs with larger vector distances in both IT_COR_ and V1_COR_.

Figure 5c (left) shows the results of the GLME analysis of the single-participant, single-trial RTs. The vector distances significantly predicted RT for both IT_COR_ (r^2^ = .23), and V1_COR_ (r^2^ = .16). When the V1_COR_ and IT_COR_ distances were entered as joint predictors in a single GLME analysis (Figure 5c, right), the IT_COR_ distances explained significant unique variance (r^2^ = .09), but the V1_COR_ distances did not (r^2^ = .02; see Table 4 for test statistics). A full model including the vector distances from all four areas of CORnet predicted the single-trial RTs moderately well (r^2^ = .36, χ^2^(8) = 33.89, p < .0001).

**Table 4.** Test statistics for analyses of single-trial response times (Experiments 3)

|  | **Experiment 3 (Adjusted)** | **Experiment 3 (Unadjusted)** |
| --- | --- | --- |
| Unique Contribution of Linear $V1_{COR}$ | $F(1,2457) = 33.43, p < .0001^*$ | $F(1,2457) = 10.04, p = .0016^*$ |
| Unique Contribution of Quadratic $V1_{COR}$ | $F(1,2457) = 2.31, p = .1285$ | $F(1,2457) = 0.27, p = .6045$ |
| Unique Contribution of Linear $IT_{COR}$ | $F(1,2457) = 52.02, p < .0001^*$ | $F(1,2457) = 17.08, p < .0001^*$ |
| Unique Contribution of Quadratic $IT_{COR}$ | $F(1,2457) = 2.04, p = .1533$ | $F(1,2457) = 3.01, p = .0828$ |
| Unique Contribution of Linear Similarity | $F(1,2457) = 175.75, p < .0001^*$ | |
| Unique Contribution of Quadratic Similarity | $F(1,2457) = 29.56, p < .0001^*$ | |
| Unique Contribution of Linear $V1_{COR}$ | $F(1,2455) = .47, p = .4913$ | $F(1,2455) = 1.42, p = .2329$ |
| Unique Contribution of Linear $IT_{COR}$ | $F(1,2455) = 12.65, p = .0004^*$ | $F(1,2455) = 5.75, p = .0166^*$ |
| Unique Contribution of Quadratic $V1_{COR}$ | $F(1,2455) = 1.04, p = .3091$ | $F(1,2455) = 0.13, p = .7147$ |
| Unique Contribution of Quadratic $IT_{COR}$ | $F(1,2455) = 0.32, p = .5694$ | $F(1,2455) = 1.01, p = .3150$ |
| Unique Contribution of Adjusted Linear $V1_{COR}$ | | $F(1,2455) = 15.32, p < .0001^*$ |
| Unique Contribution of Unadjusted Linear $V1_{COR}$ | | $F(1,2455) = 0.32, p = .5707$ |
| Unique Contribution of Quadratic Adjusted $V1_{COR}$ | | $F(1,2455) = 1.85, p = .1741$ |
| Unique Contribution of Quadratic Unadjusted $V1_{COR}$ | | $F(1,2455) = 0.34, p = .5607$ |
| Unique Contribution of Adjusted Linear $IT_{COR}$ | | $F(1,2455) = 33.51, p < .0001^*$ |
| Unique Contribution of Unadjusted Linear $IT_{COR}$ | | $F(1,2455) = 1.00, p = .3171$ |
| Unique Contribution of Quadratic Adjusted $IT_{COR}$ | | $F(1,2455) = 0.25, p = .6194$ |
| Unique Contribution of Quadratic Unadjusted $IT_{COR}$ | | $F(1,2455) = 0.07, p = .7900$ |

### Effects of adjusting for center bias

The model’s predictive ability was weaker for the focal changes to a single object in Experiment 3 than for the broadly distributed changes in Experiments 1 and 2. We anticipated this prior to seeing the results, because the architecture of CORnet does not explicitly account for the overrepresentation of the center of gaze that characterizes human visual cortex and, therefore, does not directly account for the fact that the changed items varied in eccentricity across scene pairs. After seeing that the model’s predictive ability was indeed weaker in this experiment, we created a new version of the model in which the vectors were computed after adjusting for distance from the image center. To implement this *center-bias* adjustment, the center-bias weights from the GBVS package (Harel et al., 2006) were applied to all color channels of the input image via pointwise multiplication prior to passing the image through the network. The vector distances were then computed from these adjusted images.

As shown in Figures 5d and 5e (left), the adjusted vector distances yielded a strong effect for IT_COR_ (r^2^ = .46) and a somewhat weaker effect for V1_COR_ (r^2^ = 0.31). We directly compared the adjusted and unadjusted vectors in the same GLME analysis, and we found that the adjusted vectors accounted for significant unique variance (Adjusted V1_COR_: r^2^ = .16; Adjusted IT_COR_: r^2^ = .25), but the unadjusted vectors did not (r^2^ < .01; see Table 4 for test statistics). These results show the importance of accounting for the overrepresentation of the center of gaze.

When we included the adjusted vector distances for both V1_COR_ and IT_COR_ in a joint analysis (Figure 5e, right), we found that the adjusted IT_COR_ vector distances accounted for significant unique variance in the change detection RTs (r^2^ = .16), but the adjusted V1_COR_ vector distances did not (r^2^ = .01) A full model including the adjusted vector distances from all four areas of CORnet accounted for 55% of the variance across scene pairs in the single-trial RTs (r^2^ = .55, χ^2^(8) = 58.53, p < .0001).

### Ability of RGB bitmaps to predict change detection performance

We computed vector distances from the RGB bitmaps to provide a lower reference point for variance explained. When this was done without adjusting for center bias, we found that the RGB vector distances accounted for effectively 0% of the item-level variance (χ^2^(2) = 0.04, p = .980). When we applied the center bias adjustment prior to computing the vector distances, we found that the RGB vector distances accounted for 5% of the item-level variance, but this effect was still not statistically significant (χ^2^(2) = 3.66, p = .160). Thus, whether or not we applied an adjustment to account for the eccentricity of the changed item, our model of working memory explained substantially more item-level variance in change detection RT than did the simple RGB bitmap representations of the scenes.

### Ability of similarity ratings to predict change detection performance

We next asked how well the ratings of perceptual similarity for each scene pair predicted the change detection RTs. The intraclass correlation across observers was excellent (r(87, 696) = .940, p < .0001). We found that the ratings could account for 75% of the item-level variance (r^2^ = .75, χ^2^(2) = 99.60, p < .0001). This is substantially higher than the 36% of variance explained by the full model of working memory without adjusting for center bias, but only moderately higher than the 55% of variance explained by our model after adjusting for center bias.

### Unique predictive ability of the model versus the similarity ratings

We also asked whether our model could account for variance in change detection behavior that was not captured by the similarity ratings. As in Experiment 1, this was a high bar given that higher similarity ratings were associated with smaller vector distances for both V1_COR_ and IT_COR_ (center-bias adjusted; r(78) = -0.339, p = 0.002 and r(78)= -0.445, p <.001). We therefore focused on the full model containing all four areas of CORnet with the adjustment for center bias. As in Experiment 1, we found that this full model explained significant variance in RT after accounting for perceived similarity (r^2^ = .10, χ^2^(8) = 130.67, p < .0001). In addition, the similarity ratings accounted for substantial variance in RT after controlling for the full CORnet model (r^2^ = .29, χ^2^(2) = 72.12, p < .0001).

When we included only V1_COR_ and IT_COR_ (center-bias adjusted) in the analysis, we found that this two-area model still explained 8% of the variance after controlling for the similarity ratings (χ^2^(4) = 23.10, p < .0001). In addition, the similarity ratings accounted for significant variance after controlling for the two-area model (r^2^ = .35, χ^2^(2) = 75.06, p < .0001). The unique variance explained by V1_COR_ and IT_COR_ in isolation after controlling for the similarity ratings is provided in supplementary Table S1.

## Effect of Normalization across all three Experiments

The present model involves applying a normalization operation when computing the distance between the vectors for the two images in a pair. To determine whether this normalization operation had much impact, we first asked whether the normalized vector distances differed much from the vector distances computed without any normalization. We found that the set of unnormalized distances for the scene pairs in this experiment was highly correlated with the set of normalized distances in every area of CORnet for all three experiments (*r* = .821–.998; see Table 5), especially after the adjustment for center bias in Experiment 3 (*r* = .821–.994 pre-adjustment and *r* = .926–.998 post-adjustment).

**Table 5.**
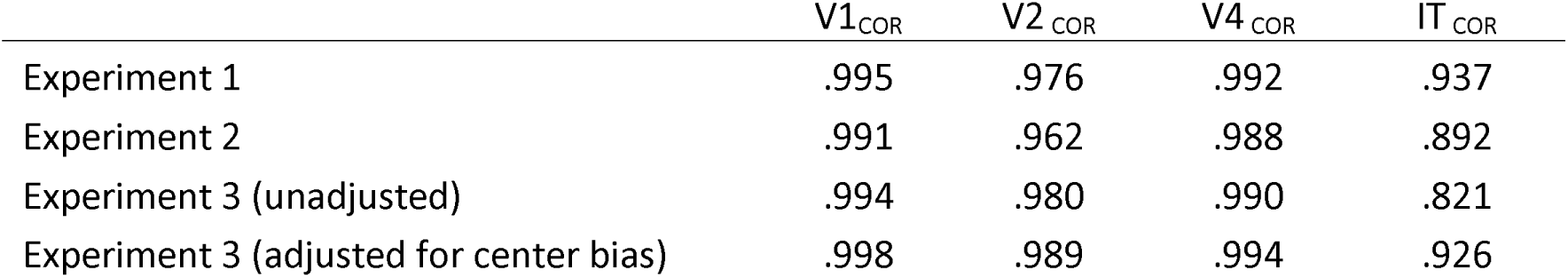
Correlation (Pearson r) between the unnormalized and normalized vector distances for each area of CORnet in Experiments 1-3.

|  | V1<sub>COR</sub> | V2<sub>COR</sub> | V4<sub>COR</sub> | IT<sub>COR</sub> |
| --- | --- | --- | --- | --- |
| Experiment 1 | .995 | .976 | .992 | .937 |
| Experiment 2 | .991 | .962 | .988 | .892 |
| Experiment 3 (unadjusted) | .994 | .980 | .990 | .821 |
| Experiment 3 (adjusted for center bias) | .998 | .989 | .994 | .926 |

We followed this analysis by repeating the main statistical analyses from the previous section for each experiment, except using the unnormalized vector distances rather than the normalized vector distances to predict the change detection RTs. The pattern of results (shown in supplementary Figures S2 and S3) was very similar to the pattern observed for the normalized vector distances (Figures 3 and 5). The r^2^ values for these results are shown in Table 6, with the test statistics corresponding to these values in Supplementary Table S4, S5 and S6. Generally, the impact of normalization was minor and inconsistent, with the r^2^ values for two approaches being within .05 of each other in all cases. Thus, in the specific image sets used in all these experiments, normalization played only a minor role in the ability of the model to predict behavior.

**Table 6.** r^2^ values for the unnormalized and normalized vector models for Experiments 1-3.

|  | V1<sub>COR</sub> | IT<sub>COR</sub> | Unique<br>V1<sub>COR</sub>/IT<sub>COR</sub> | Full Model |
| --- | --- | --- | --- | --- |
| Experiment 1 normalized | .44 | .57 | .02/.15 | .76 |
| Experiment 1 unnormalized | .48 | .57 | .04/.19 | .71 |
| Experiment 2 normalized | .36 | .71 | <.01/.35 | .80 |
| Experiment 2 unnormalized | .39 | .61 | <.01/.36 | .76 |
| Experiment 3 normalized (unadjusted) | .16 | .23 | .02/.09 | .36 |
| Experiment 3 unnormalized (unadjusted) | .14 | .32 | .01/.22 | .41 |
| Experiment 3 normalized (CB adjusted) | .31 | .46 | .01/.16 | .55 |
| Experiment 3 unnormalized (CB adjusted) | .32 | .47 | .02/.24 | .57 |

## Experiment 4: Patterns of Neural Activity During the Delay Period

A model of working memory should go beyond predicting behavioral performance at the time of decision, which might be influenced by a variety of factors beyond the working memory representations (Yoo et al., 2021). The goal of Experiment 4 was therefore to test whether the population vector model can also account for the patterns of neural activity during the delay period, while the working memory representation is being maintained. Brain activity was measured via EEG, which has the temporal resolution necessary to isolate activity during the short delay periods that are typically used to isolate working memory (Jeneson et al., 2012). Representational similarity analysis (RSA; Kriegeskorte et al., 2008) was used to link brain activity to the model.

Whereas Experiments 1-3 used the distance between the population vectors for the two images on a given trial to predict change detection performance, only one image was presented on each trial in Experiment 4, and we tested whether the vector for this image was related to the pattern of voltage across the scalp at each time point during the delay period.

Because the EEG is a very noisy signal, we repeated each of the images multiple times so that we could compute an averaged waveform for each image (i.e., an *event-related potential* or *ERP* waveform). To obtain a sufficient number of trials, we used the modified 1-back task shown in Figure 6a. In this task, memory was tested on only 10% of trials (to minimize the amount of time spent obtaining behavioral responses). To motivate participants to form a detailed memory on every trial, we tested their memory by showing them a small region of the immediately preceding image and requiring them to report whether this region was in the original location or shifted slightly. A staircase procedure was used to ensure a consistent level of task difficulty across participants and across trials. Note that this task was optimized for the EEG analyses, and it was not designed to produce behavioral data that could be used to test the model (especially given that the size of the shift varied over trials).

**Figure 6.**
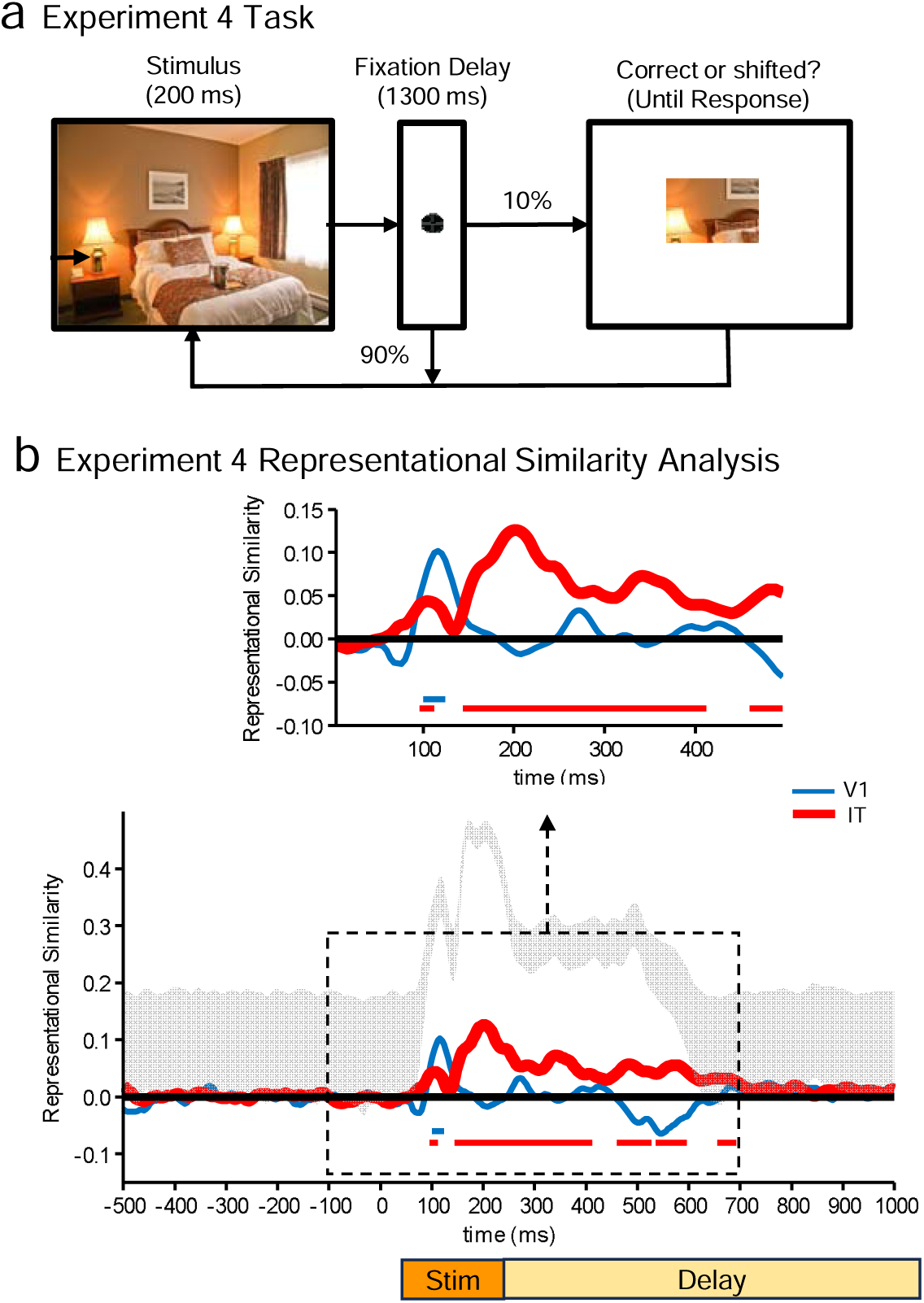
Experiment 4 Task and Results. *Note.* **a**, Task. Participants saw a sequence of scenes, each followed by a delay. For 10% of stimuli, the delay was followed by a snippet of the scene, and the participant was required to report whether this snippet was in the original location or was shifted diagonally. The size of the shift was continually adjusted to maintain a constant level of performance. **b**, Representational similarity between the V1_COR_ or IT_COR_ vectors and the pattern of voltage over the electrodes, computed separately at each time point. The bottom portion shows the upper and lower bounds of the estimated noise ceiling of the ERP data as the gray shaded region, and the upper portion shows the key time region without the noise ceiling so that the waveforms can be seen more clearly. Horizontal line segments across the bottom indicate time periods in which the representational similarity values are significantly greater than zero (p < .05) after a false discovery rate correction for multiple comparisons.

## Method

### Participants

The participants in this study were adults recruited from the University of California, Davis community. They were paid $15/hour for participating. We tested 31 participants in total, 2 of whom were excluded because of artifacts, leaving a final sample of 29 participants (see Table 1 for demographic information).

The target sample size for this experiment was chosen a priori based on an empirical power analysis in which we simulated results under the design parameters of the current study with varying sample sizes. On the basis of pilot data, we used a representational similarity effect size of 0.10 (*SD* = .05) over a 100-ms period. The simulation results showed that a sample size of 10 was sufficient to detect significant effects for 80% of the time points within the analysis window after correction for multiple comparisons (see below for details). However, we targeted an *N* of 30 to provide more precise parameter estimates.

### Stimuli

The stimuli were presented using Matlab with PsychToolbox (Brainard, 1997; Pelli, 1997) on an HP ZR2440W LCD monitor with a gray background at a viewing distance of 100 cm. The images consisted of 50 photographs of real-world scenes that had been used in previous research on scene processing (Henderson & Hayes, 2017, 2018; Kiat, Hayes, et al., 2022), each subtending 8° × 6° of visual angle. None of the scenes included humans or animals. The images and experimental control script are available at https://osf.io/7ukgj/.

### Procedure

As shown in Figure 6a, the image for a given trial was presented for 200 ms, followed by a 1300-ms delay period. On 90% of trials, the next image began immediately at the end of the preceding delay period. For the remaining 10% of trials, memory for the image on that trial was probed at the end of the delay period. In a probe display, a randomly selected 3° × 2° portion of the scene was shown, either presented in the original location or shifted diagonally by a small amount (with equal probability). Participants used a game controller to report whether this portion of the scene was presented at the original location or shifted (using the left index finger to indicate a shift and the right index finger to indicate no shift). Accuracy rather than speed was emphasized. The shift amount varied across trials according to a 3-down/1-up staircase, leading to approximately 80% correct responses. This ensured that task difficulty was approximately equal across participants and that it remained relatively constant over the course of the session.

Each of the 50 images was shown a total of 32 times. The images were shown in random order in 16 blocks, with each block consisting of one presentation of each of the 50 scenes in random order. A practice block of 50 trials using a different set of scenes preceded the main task.

### EEG recording and preprocessing

Electrophysiological activity was recorded using a Brain Products ActiCHamp recording system with a sampling rate of 500 Hz and a cascaded integrator-comb antialiasing filter (half-power cutoff at 130 Hz). We recorded the EEG from 59 scalp sites (AF3, AF4, AF7, AF8, FC1, FC2, FC3, FC4, FC5, FC6, FP1, FP2, F1, F2, F3, F4, F5, F6, F7, F8, C1, C2, C3, C4, C5, C6, CP1, CP2, CP3, CP4, CP5, CP6, P1, P2, P3, P4, P5, P6, P7, P8, P9, P10, PO3, PO4, PO7, PO8, T7, TP7, TP8, O1, O2, Fz, FCz, Cz, CPz, Pz, POz, and Oz), from the left and right mastoids, and from three electrooculogram (EOG) sites. Horizontal EOG electrodes were placed lateral to the left and right external canthi, and a vertical EOG electrode was placed below the right eye. Electrode impedances were maintained at <15 kΩ. All signals were recorded single-ended and then referenced offline. See Farrens et al. (2019) for a detailed description of the electrode application and recording protocol.

All preprocessing steps were conducted in MATLAB using the EEGLAB (Delorme & Makeig, 2004) and ERPLAB (Lopez-Calderon & Luck, 2014) toolboxes following a standard pipeline (Luck, 2022). The processing scripts and data are provided at https://osf.io/7ukgj/. First, the event codes were shifted to account for monitor delay (24 ms as measured by a photodiode). The EEG signals were then referenced to the P8 electrode. A bipolar horizontal EOG derivation was then computed as the difference between the two horizontal EOG electrodes, with a vertical EOG derivation computed as the difference between FP2 and the vertical EOG electrode. All the signals were then bandpass filtered (noncausal Butterworth impulse response function, DC offset removed, half-amplitude cutoffs at 0.1 and 20 Hz, 12 dB/oct roll-off) and resampled at 250 Hz (one sample every 4 ms). Time sections between blocks were then deleted from the datasets.

Independent component analysis (ICA) was then performed using the Infomax algorithm on the retained continuous EEG for each participant to identify and remove components that were associated with blinks (Jung et al., 2000) and eye movements (Drisdelle et al., 2017). We used an optimized procedure (Luck, 2022) in which a copy of the original data was filtered more aggressively (noncausal Butterworth impulse response function, half-amplitude cutoffs at 1 – 20 Hz, 48 dB/oct roll-off) prior to the ICA decomposition to better separate ocular artifacts from neural signals. The ICA weights derived from this decomposition were then transferred back to the original data and used to reconstruct the EEG data without independent components corresponding to blinks and eye movements (typically 1-2 independent components per participant). The criterion for excluding an ICA component from the reconstruction was the consistency between the shape, timing, and spatial location of the component compared with the bipolar HEOG and VEOG signals.

The ICA-corrected EEG signals were then segmented for each trial from -500 to +1500 ms relative to the onset of the scenes and baseline-corrected using the mean voltage from -500 to 0 ms. Individual trials were then rejected if the peak-to-peak voltage was >200 μV in any 200 ms window in any electrode, or if a blink or eye movement (defined as a step-like voltage change; Luck, 2014) was detected in the uncorrected bipolar HEOG or VEOG signals between 200 ms prestimulus and 200 ms post-stimulus (and might therefore impact the perception of the stimulus, which ended at 200 ms). We always exclude participants for whom more than 25% of trials were rejected; two of the 31 participants were excluded from the final sample for this reason. An average of 4.8% (SD = 4.9%) of trials were rejected in the remaining 29 participants.

For each participant, we created an averaged ERP for each of the 50 scenes by averaging together all the unrejected single-trial EEG segments. Supplementary Figure S4 shows the grand average waveform across participants and scenes at representative electrode sites, along with the mean global field power across sites (Skrandies, 1989).

### Representational Similarity Analysis (RSA)

We used RSA to link the neural signals from the averaged ERPs for each individual scene to the activation vectors produced by those scenes in a given area of CORnet-S. RSA has the advantage of abstracting the data away from the original units and into the underlying representational geometry. It is performed independently for each participant and can thus tolerate differences in overall scalp voltage patterns across participants resulting from differences in cortical folding patterns. In addition, it is performed independently at each time point (every 4 ms), yielding excellent temporal resolution. Despite the relatively low spatial resolution of EEG signals, RSA works reasonably well with averaged ERP data (Greene & Hansen, 2018; T. He et al., 2022; Kaiser & Nyga, 2020; Kiat, Hayes, et al., 2022; Robinson et al., 2017; Salmela et al., 2018).

When applied to ERPs, RSA involves quantifying the similarity between the neural responses to each pair of stimuli. To accomplish this, we first extracted the pattern of voltages across the 59 electrode sites at a given time point in the averaged ERP waveform for each of the 50 scenes. We used the Pearson *r* correlation between the set of voltages across electrode sites for one scene and the set of voltages across electrode sites for another scene. We computed this *r* value for each of the 50 x 50 pairs of scenes, creating a 50 × 50 *representational similarity matrix* (RSM) for the neural data. (Other similarity metrics are possible, but we find that the Pearson *r* works particularly well in ERP experiments like this.) A separate RSM was computed at each time point (every 4 ms from -500 ms to 1500 ms relative to scene onset) for each participant.

We also quantified the similarity between each pair of scenes in terms of the pattern of activation across the units within a given area of CORnet-S. That is, we fed each scene into the network, obtained the vector of activation values across the ReLU units of the final layer of a given area, and computed the Pearson *r* correlation between the vectors for each pair of scenes. Note that we intentionally used the same similarity metric (Pearson *r*) for both the neural data and the model. Because we had 50 scenes, this process yielded a 50 × 50 RSM for each area of CORnet.

To estimate the representational similarity between the neural data and the computational model, we computed the Spearman rank-order correlation between the neural RSM at a given time point and the CORnet RSM for a given area (using only the lower triangle of values in each RSM because the upper and lower triangles contain identical values). This yielded a waveform of representational similarity values, one per time point, for each area of CORnet, seperately for each participant. We then ran statistical analyses to determine the time points at which the mean across participants of the representational similarity values was significantly greater than zero. Separate analyses were conducted for each combination of ERP time point and CORnet area.

Our statistical analysis approach was identical to that used in all of our previous RSA studies (e.g., T. He et al., 2022; Kiat, Hayes, et al., 2022; Kiat, Luck, et al., 2022). First, we used a nonparametric Wilcoxon signed-rank tests to avoid making parametric assumptions about the data (which deals with the non-normality of the Spearman rho correlations used to quantify representational similarity). Second, negative RSA values are typically meaningless and likely to be spurious in this type of research, so we did not try to interpret negative values, and we used one-tailed tests of significance. Third, we applied the false discovery rate (FDR) correction for multiple comparisons (Benjamini & Hochberg, 1995) for all analyses, using a testing period of 100 to 1000 ms poststimulus. Note that this method of correction avoids several downsides of cluster-based correction methods (see, e.g., Sassenhagen & Draschkow, 2019).

The main analyses presented here were conducted separately for V1_COR_ and IT_COR_. To parallel the behavioral analyses, we also conducted a rank regression procedure in which the ERP RSMs were jointly regressed onto the V1_COR_ and IT_COR_ RSMs, allowing us to isolate the unique variance accounted for in the ERP data by each individual area. In other words, variance in the V1_COR_ RSM that was explained by the IT_COR_ RSM was partialled out when examining the correlation between the ERP RSM and the V1_COR_ RSM, and vice versa.

We also estimated the *noise ceiling* at each time point, which indicates the maximum representational similarity that could be expected given the noise level of the ERP data at each time point. More concretely, the noise ceiling quantifies the consistency of the ERP RSMs across participants. Following a standard approach (Nili et al., 2014), the upper bound of the noise ceiling was estimated by computing the Pearson *r* correlation between the RSM for one participant and the grand average RSM across all participants, and then averaging the *r* values that were obtained for each participant. The lower bound of the noise ceiling was estimated in the same manner, except that the grand average RSM used to compute the *r* value for a given participant excluded that participant. All of the RSA scripts, including those for computing the noise ceiling, are available at https://osf.io/7ukgj/.

## Results and Discussion

Figure 6b shows the representational similarity between the neural signals at each time point and the V1_COR_ and IT_COR_ population vectors (results for all four areas are shown in supplementary Figure S5). For V1_COR_, we observed strong representational similarity between the model and the ERPs only during the sensory processing period. The representational similarity was significantly greater than zero for V1_COR_ from 100-132 ms. We also observed an initial peak for IT_COR_ during sensory processing, which was significantly greater than zero from 100-124 ms. Unlike V1_COR_, we also observed sustained representational similarity for IT_COR_ during the delay period (significantly greater than zero from 144-716 ms). These results demonstrate that our model can account for differences in the patterns of brain activity produced by different scenes while the scenes are being maintained in working memory.

As time passes from the baseline, low-frequency contamination of the EEG from skin potentials and other sources causes a gradual buildup of noise in the ERP waveform (Luck, 2014; G. Zhang & Luck, 2023). This noise buildup will cause representational similarity to become attenuated at longer latencies (all else being equal), which might explain the drop in representational similarity observed after approximately 700 ms in the present study. The computed noise ceiling, which indicates the maximum representational similarity that could be expected at each time point given the noise, provides some insights into this. As shown in Figure 6b, the noise ceiling dropped sharply from approximately 500–700 ms, which necessitated a drop in representational similarity during this time period. Thus, the lack of sustained representational similarity after approximately 700 ms likely reflects technical limitations rather than a change in the underlying neural activity.

The results shown in Figure 6b reflect independent analyses of V1_COR_ and IT_COR_. This approach was justified because the rank correlation between the V1_COR_ and IT_COR_ RSMs was fairly low for the images used in this experiment (*r* = .127, *p* = .0199). Supplementary Figure S6 shows that the pattern of results was nearly identical when we used partial correlations to isolate the unique contribution of the V1_COR_ and IT_COR_ areas, indicating that the predictive contribution of each area in this analysis was fairly independent.

Note that negative representational similarity values were observed from approximately 80-100 ms for V1_COR_. Negative values were also observed across a wide range of time points for V2_COR_ (see supplementary Figure S5). As noted earlier, negative representational similarity values such as these are not typically interpretable and are likely to be spurious. That is, although one can easily create artificial representational similarity matrices that produce negative representational similarity values, it is difficult to conceive of realistic scenarios in which this would occur with actual neural data. For example, inverting the ERP voltage values produced by each scene would not change the scene-to-scene similarity values in the representational similarity matrices. Consequently, non-spurious representational similarity values are almost always positive (Schütt et al., 2023).

## General Discussion

Visual working memory plays an essential role in everyday visually guided behavior, but almost all formal models of visual working memory have focused on arrays of discrete, simple, artificial objects such as colored squares (Bays, 2014; Rouder et al., 2008; Schurgin et al., 2020; van den Berg et al., 2014; W. Zhang & Luck, 2008). These models fail to capture essential aspects of real-world scenes, such as irregular shape contours and color gradients (Luck & Kiat, 2024b). Here, we have developed a quantitative model of working memory based on the general idea that the working memory representation of a scene is a weak/noisy version of the perceptual representation of the scene encoded by the vector of firing rates across a population of neurons in one or more regions of the ventral visual pathway.

We do not have access to the actual vector of firing rates, so we implemented this theory in a formal model using the vector of activation values produced by each scene in CORnet-S, a DNN that was designed to reflect the known anatomical properties and activation patterns of the ventral visual pathway (Kubilius et al., 2018). Although CORnet-S is not a perfect model of the ventral pathway, it is the best available model, and it provides a practical solution for implementing our theory of working memory as a formal model while we await better models of the ventral stream.

This model of working memory passed several tests, including the ability to predict behavioral performance very well and the ability to account for differences in patterns of brain activity for different scenes. These results establish the viability of the population vector model of working memory. Moreover, the extensive data and code that we have made freely available can be used to pit our model against other quantitative models of working memory for natural scenes once such models exist.

The following sections will summarize the evidence supporting the viability of the model along with the limitations of the model in its current form.

### Comparison with vectors from the RGB bitmaps

The population vector model as implemented here predicted behavioral change detection performance remarkably well. It dramatically outperformed a simplistic model in which vector distances were obtained directly from the RGB bitmaps of the image pairs. These bitmaps contain all of the information needed to perform the change detection task, but in a raw format that is unlikely to be similar to the format of working memory representations, which classic research has shown are more abstract than bitmaps (Phillips, 1974; Sperling, 1960). The fact that the bitmaps contain all the information needed to perform the task—but in a format that is unlikely to reflect working memory—made them a good minimum reference point for comparison. The finding that our model dramatically outperformed this reference point was an important first step in demonstrating the viability of the model. Note also that the poor performance of the RGB bitmaps, especially in Experiments 3 and 4, rules out the possibility that vectors from any information-rich representations of the scenes would be able to explain behavioral performance and patterns of neural activity. Thus, to account for the behavioral data and the EEG data, it is necessary to have a model that transforms the input into an appropriately abstract representation.

### Comparison with perceptual similarity ratings

We also compared the predictive ability of our model with the predictive ability of perceptual similarity ratings of the scene pairs in Experiments 1 and 3^3^. The perceived similarity of a given pair of scenes when presented simultaneously in the rating task should do an excellent job of predicting how quickly and accurately people could judge whether those same pairs were the same or different when presented sequentially in our change detection tasks. That is, the more similar the scene pairs appear to be when presented side by side, the more difficult it should be to detect a difference between them when presented sequentially. We therefore used the predictive ability of these similarity ratings to provide an upper reference point for the predictive ability of our model. Specifically, we assumed that if the amount of variation in change detection performance across scene pairs explained by our model comes anywhere near the amount of variation explained by the perceptual similarity ratings, then this would indicate that our model is viable and worthy of continued development.

Indeed, we found that our model (including all four CORnet areas) accounted for 76% of the item-level variance across scene pairs in Experiment 1, which was almost as much as the 78% accounted for by the ratings. In Experiment 3, we found that our model (after adjustment for center bias) accounted for 55% of the item-level variance in change detection performance, which compared well with the 75% of variance explained by the similarity ratings. Although we could not realistically obtain similarity ratings for Experiment 2, the model accounted for an impressive 80% of the item-level variance in that experiment.

Thus, the population vector model predicted both the speed and accuracy of change detection performance extremely well, much better than the lower reference point provided by the RGB bitmaps and nearly as well as the perceptual similarity ratings. The model even explained a small amount of unique variance that was not explained by the similarity ratings. This may indicate that observers weigh the various image features differently when making an explicit judgment of similarity for side-by-side images than when making a same-different judgment for sequential images, leaving some variance to be explained by the model. However, we cannot rule out the possibility that this instead reflects some imperfection in the similarity ratings (e.g., resulting from noise or from the nature of the rating task), leaving some variance to be explained by the model. However, even if a perfect set of ratings could explain 99% of the variance in behavior, leaving no residual variance to be explained by the model, the finding that our model can explain as much as 80% of the item-level variance in behavior indicates that the model is viable and worthy of further development.

### Neural activity during the delay period

The tasks used in Experiments 1-3 required participants to hold one stimulus in memory and then compare this working memory representation with the perceptual representation of the next stimulus. It is possible that our model’s ability to predict the speed and accuracy of change detection in those experiments was influenced by the operations and representations involved in perceiving the second stimulus in a pair, comparing it with the working memory representation of the first stimulus, and then deciding whether to report that a change was present. Because our model is supposed to specify the nature of the working memory representation that is maintained during the delay between one stimulus and the next, it is vital to assess the model’s ability to account for the nature of the delay-period representation without the potentially confounding influences of the comparison process. To assess this aspect of the model, we used representational similarity analysis to examine whether image pairs that produced similar spatial patterns of EEG over the scalp during the delay period also produced similar activity vectors in the model. As predicted, we found significant representational similarity between the EEG signals and the model during both the perceptual processing interval and the delay period (especially for the IT_COR_ area). By contrast, vectors from the RGB bitmaps exhibited no significant representational similarity with the EEG activity during the delay period. These results provide important additional support for the viability of the population vector model.

### Capacity limits

The population vector model does not explain limitations in memory performance using concepts such as “items” or “resources.” Indeed, it is designed to operate on scenes that are not easily parsed into simple, discrete items. According to our model, performance naturally degrades to the extent that the spatial architecture and feature channels of the network yield similar representations for the two scenes in a pair. For example, the units in IT_COR_ (like neurons in primate inferotemporal cortex) have large receptive fields that integrate information across a broad spatial range, which will make it difficult for IT_COR_ to distinguish between scenes that differ slightly in the locations of the scene elements. Thus, the fact that the model correctly predicts poor performance for some scene pairs and good performance for others is likely a result of the architecture and feature tuning of the model.

In addition, the population vector model includes a normalization operation that was inspired by Weber’s law (Fechner, 1948), and this may also produce something akin to capacity limitations. Specifically, the ability of the model to distinguish between two versions of a scene is scaled by the vector length of the scenes, which means that the model’s ability to detect a change of a given absolute magnitude will be poorer for complex scenes (which strongly activate a large number of neurons and produce a long vector length) than for simple scenes (which activate fewer neurons and produce a short vector length).

However, it turns out that the normalization operation played relatively little role in the present results. For the images used in these experiments, the normalization operation had a similar impact on all the scenes, and the normalized vector distances were very highly correlated with the unnormalized vector distances. Moreover, the ability of the model to predict behavior was comparable with and without normalization. However, normalization may be important for predicting performance in experiments using a broader range of scenes. This is an open question for future research.

The normalization operation in the present model broadly resembles the divisive normalization operation used to explain capacity limitations in some models of working memory for simple objects (Bays, 2014; Bouchacourt & Buschman, 2019; Wei et al., 2012). However, whereas the normalization in those other models is essential for the models to account for differences in behavioral performance across different stimuli (e.g., across set sizes), the normalization in the present model had relatively little effect on the ability to predict differences in performance across different stimuli (i.e., scene pairs) in the present experiments.

### Precision, individual differences, and pattern separation

Although the present model does not contain any direct limits on storage capacity (as defined in terms of items), the related concept of *precision* can be directly mapped onto the model. A representation is precise to the extent that it is the same each time a given input is presented (i.e., high precision is equivalent to low trial-to-trial variability). Precise representations therefore make it possible to reliably distinguish between highly similar input patterns. In the population vector model, trial-to-trial variability in the representations occurs as a result of noise in the activation of the units. Consequently, a participant with less noise (more precise representations) would exhibit better performance in the tasks used here than a participant with more noise (less precise representations).

The current version of the population vector model does not account for individual differences in noise or precision. This is a limitation of the present approach, because modeling the noise level for each participant would make it possible to quantify individual differences in working memory, which can be quite valuable (Engle et al., 1999; Unsworth & Engle, 2007; Vogel & Awh, 2008). However, it should be possible to develop a procedure to model the noise level for individual participants in the context of the population vector model and then ask whether individual differences in noise are correlated with individual differences in other variables. This will be an important direction for future research.

The concept of pattern separation is also relevant here (see, e.g., Bakker et al., 2008; Yassa & Stark, 2011). Specifically, the medial temporal lobes contain circuitry that increases the separability of representations of similar input patterns. In the context of the present model, such circuitry would enhance behavioral performance for scene pairs that are highly similar. Note that, although the medial temporal lobes are traditionally associated with long-term memory (see, e.g., Jeneson et al., 2012), recent evidence suggests that they also play a key role in working memory tasks that require high levels of precision (see, e.g., Borders et al., 2022; W. Xie et al., 2023).

### Combining with a broader model of memory

It may be possible to gain greater insights into these issues by combining the population vector model with the target confusability competition (TCC) model of Schurgin et al. (2020). When applied to natural scenes, the TCC model would propose that the activation of a to-be-remembered sample scene in memory would cause a spreading of activation to the memory representations of other scenes as a function of their similarity to the sample scene. These activation levels are then used to make a decision when a test scene is presented. Because of noise, the actual activation level for a given test scene will vary across trials even if the sample scene is held constant. If the noise level is increased, this increases the likelihood that the activation level for a given test scene will be greater than the activation level for the original sample scene, which may lead to an incorrect behavioral response. Thus, behavioral performance for discriminating between a given sample scene and a given test scene will depend on both the intrinsic similarity between the two scenes and the amount of noise. In the population vector model, the intrinsic similarity between two scenes is simply the length of the vector between the representations of the two scenes in a given area of cortex. Bates et al. (2024) have already demonstrated that this general approach is feasible.

A key proposal of the TCC model is that the amount of noise can be captured by a memory strength parameter (d’): the effective difference in activation between a sample item and a test item is the intrinsic similarity between the two items multiplied by the memory strength parameter (d’). When d’ is larger, the test scene is pushed further from the sample scene, decreasing the likelihood that the test and sample scenes will be confused. Thus, memory performance can be predicted for any set of scenes as long as the intrinsic scene-to-scene similarities are known and the individual participant’s d’ is known. It should therefore be possible to estimate d’ for one set of scenes and then use the vector distances from the population vector model to predict the d’ for any other set of scenes. In principle, a separate d’ value can be estimated for each individual and used as a way of assessing individual differences. This would be an interesting direction for future research.

### Limitations of the model

The specific model presented here has a number of limitations, and it should be regarded as Population Vector Model Version 1.0. One of the most significant limitations of the current version is that it uses CORnet to estimate the population vector for each scene. CORnet has some significant virtues, but it is still far from an adequate model of the ventral stream. For example, it has no fovea, no lateral connections, and no feedback between areas. More generally, DNNs of this sort tend to overutilize texture information and make errors that are very different from human errors unless specialized training regimes are used (Baker et al., 2018; Dodge & Karam, 2017; Hermann et al., 2020). Perhaps most significantly in the present context, DNNs like CORnet provide a very poor match to neural representations for the types of artificial stimuli that have dominated research on visual working memory for the past 20 years (Xu & Vaziri-Pashkam, 2021). It is also known that current DNNs do not fully account for all explainable variance in higher ventral stream areas (Jozwik et al., 2018). Nonetheless, CORnet was a reasonable choice for version 1.0 of the model because it was the best model of the human ventral pathway that was available when we conducted the present study. The population vector model can readily be updated to use better models of the ventral stream when they become available (whether or not they are DNNs).

Because current DNNs do a poor job of accounting for representations of artificial stimuli, the current version of the population vector model is suitable only for real-world scenes. Indeed, our preliminary explorations have suggested that it cannot account for the canonical pattern of change detection results observed with arrays of artificial objects, in which performance is near ceiling for arrays of 1-2 objects and then falls rapidly as the set size increases further (Luck & Vogel, 1997; Vogel et al., 2001). Bates et al. (2024) also found that DNNs tended to do a poor job of accounting for the human pattern of set size effects for arrays of simple orientation stimuli. Similarly, Xie et al. (2023) found that a model combining a “sensory” DNN like CORnet with a “cognitive” recurrent neural network failed to exhibit the human pattern of change detection performance, in which accuracy is near ceiling for set sizes 1-3 and then falls rapidly. An adequate model of working memory should be able to explain performance for both naturalistic and artificial images, so this is a significant limitation of the current model. However, given the scarcity of formal models of working memory for real-world scenes, the present model represents a significant step forward.

Future models will also need to incorporate at least two types of attentional mechanisms. First, an *input selection* mechanism is needed to reflect the fact that people can easily choose to transfer a subset of the sensory input into working memory. For example, when instructed to do so, people can store only the information from the left side of the display or only the X-shaped items (Schmidt et al., 2002; Turvey & Kravetz, 1970; Vogel & Machizawa, 2004). This type of mechanism could presumably be implemented within the architecture of the current model by means of *block attentional modules* (Khan et al., 2020; Woo et al., 2018), which are gain-control mechanisms that control the transmission of specific features or locations from one layer to the next (much like gain control mechanisms in the neuroscience literature on attention; Hillyard et al., 1998; Itthipuripat et al., 2014; Reynolds & Heeger, 2009; Treue & Martinez Trujillo, 1999).

Second, a *decision-weighting* mechanism is needed, much like those used in models of category learning (Kruschke, 2004). Decision weighting allows a person to take an existing working memory representation and use it to perform tasks that go beyond assessing whether or not a subsequent visual input is an exact match. For example, people can easily detect a change in a shape while disregarding a change in the shape’s location (Phillips, 1974). Note that this type of mechanism does not alter what information is stored in working memory, but merely transforms the working memory representation into a behavioral response. Thus, it could be implemented by a system that reads the output of the working memory system.

### Natural scenes versus arrays of simple objects

It is possible that the addition of attention mechanisms to the population vector model—along with the use of a more advanced underlying model of the ventral stream—will allow us to accurately predict set size effects for arrays of simple artificial objects in addition to predicting differences in performance for different natural scene pairs. A different possibility, however, is that the brain contains both an item-based working memory system (like the one conceptualized by Adam et al., 2017; W. Zhang & Luck, 2008) and a scene-based working memory system (like the population vector model). The item-based system might be particularly useful when the task involves storing a high-fidelity representation of only a few items from a scene (e.g., when a person stores a representation of one object in working memory to compare one feature of this object with another object that cannot be seen at the same time). Such a system would also be very useful for laboratory tasks in which a small number of colored squares must be held in working memory. However, such a system would not be very useful when the task involves comparing two complex scenes or involves remembering complex objects that involve color gradients or continuously varying shape contours. In these cases, the scene-based working memory system might be prioritized. The question of whether and how people combine information from multiple memory signals has received considerable attention in the broader memory literature (e.g., DeCarlo, 2002; Wixted & Mickes, 2010; Yonelinas, 2024).

### Roles of V1_COR_ versus IT_COR_

There is a longstanding controversy about the roles of different visual areas in working memory (Leavitt et al., 2017; Xu, 2020, 2020). Most single-unit and fMRI studies of visual working memory have found that elevated delay-period activity is limited to mid-to-high levels of visual cortex (Bisley et al., 2004; Chelazzi et al., 1998; Fuster & Jervey, 1981; Lee et al., 2005; Miyashita & Chang, 1988; Motter, 1994). However, the *contents* of working memory can be decoded from the activation pattern across electrodes and voxels in area V1 (Ester et al., 2009; Harrison & Tong, 2009; Huang et al., 2024; Serences et al., 2009). In the present study, IT_COR_ consistently accounted for unique variance in both behavior and brain activity after controlling for V1_COR_, whereas V1_COR_ accounted for little or no unique variance after controlling for IT_COR_. The V2_COR_ results were generally similar to the V1_COR_ results, and the V4_COR_ results were generally similar to the IT_COR_ results (see supplementary Figures S1 and S5).

However, the present results cannot definitively rule out a role for human area V1 in working memory, because CORnet is an imperfect model of the visual system (e.g., it lacks lateral inhibition within areas and recurrent connections between areas). One intriguing possibility is that area V1 does represent information in working memory, but not in the same format used for perception. This would be consistent with the finding that working memory representations in V1 can be decoded for stimuli outside the receptive field (Huang et al., 2024) and even in the ipsilateral hemisphere (Ester et al., 2009; Harrison & Tong, 2009; Serences et al., 2009). Another possibility is that the activity decoded in V1 represents feedback signals from higher areas that are part of a predictive coding system (of the type proposed by Rao & Ballard, 1999).

### Constraints on Generality

The data in the present study were collected from convenience samples of neurotypical young adults living in a college town. Although there is no obvious reason why this sampling approach would lead to misleading conclusions, we cannot say from the present results whether similar findings would be obtained from other populations of participants. It would be valuable for future research to assess how these findings generalize to populations of different age ranges, cultural and educational backgrounds, and neurodivergency.

## Footnotes

1 In principle, we could implement our theory using a broad range of different DNNs and ask which DNNs led to the best predictions. However, our goal is to test our *theory*, which refers to the pattern of population activity in visual cortex, so the question of which DNN best predicts the behavioral data does not test our theory. Moreover, Bates et al. (2024) already performed a careful comparison of a broad range of DNNs.

2 We initially analyzed the data without the quadratic term. We then realized that this resulted in negative RTs being predicted for the largest vector distances, so we added the quadratic term to account for the inherent floor in RTs.

3 Note that it was impractical to obtain these ratings for Experiments 2 and 4, both of which would have required many more ratings.

