## Supplementary Materials for "A Population Vector Model of Visual Working Memory for Real-World Scenes"

**Figure S1**

*V2_COR_ & V4_COR_ Results from Experiments 1-3*

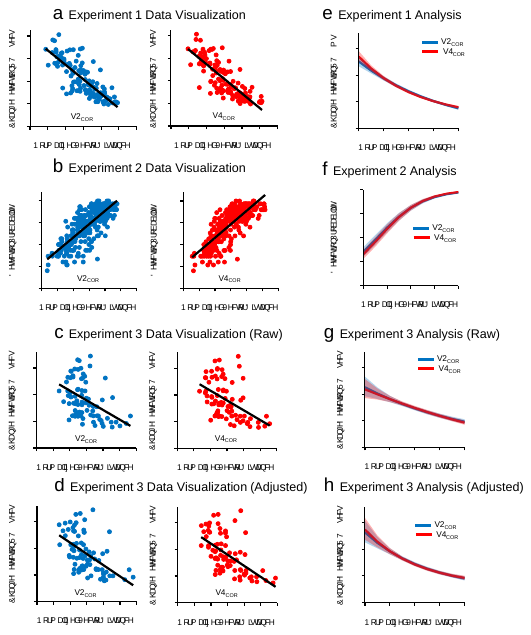

*Note.* **a,b,c,d,** Scatterplot visualization of the results from the three Experiments, showing the relationship between the V2_COR_ or V4_COR_ vector distances for a given scene pair and the mean response time (RT: Experiments 1 and 3) or change detection probability (Experiment 2) across participants for that pair. **e, f, g, h.** Results of the V2_COR_ and V4_COR_ univariate statistical models of the single-trial RTs from Experiment 1 (**e, g, h**) and the single-trial binary responses from Experiment 2 (**f**).

**Figure S2**

*Experiments 1 and 2 results using unnormalized vectors.*

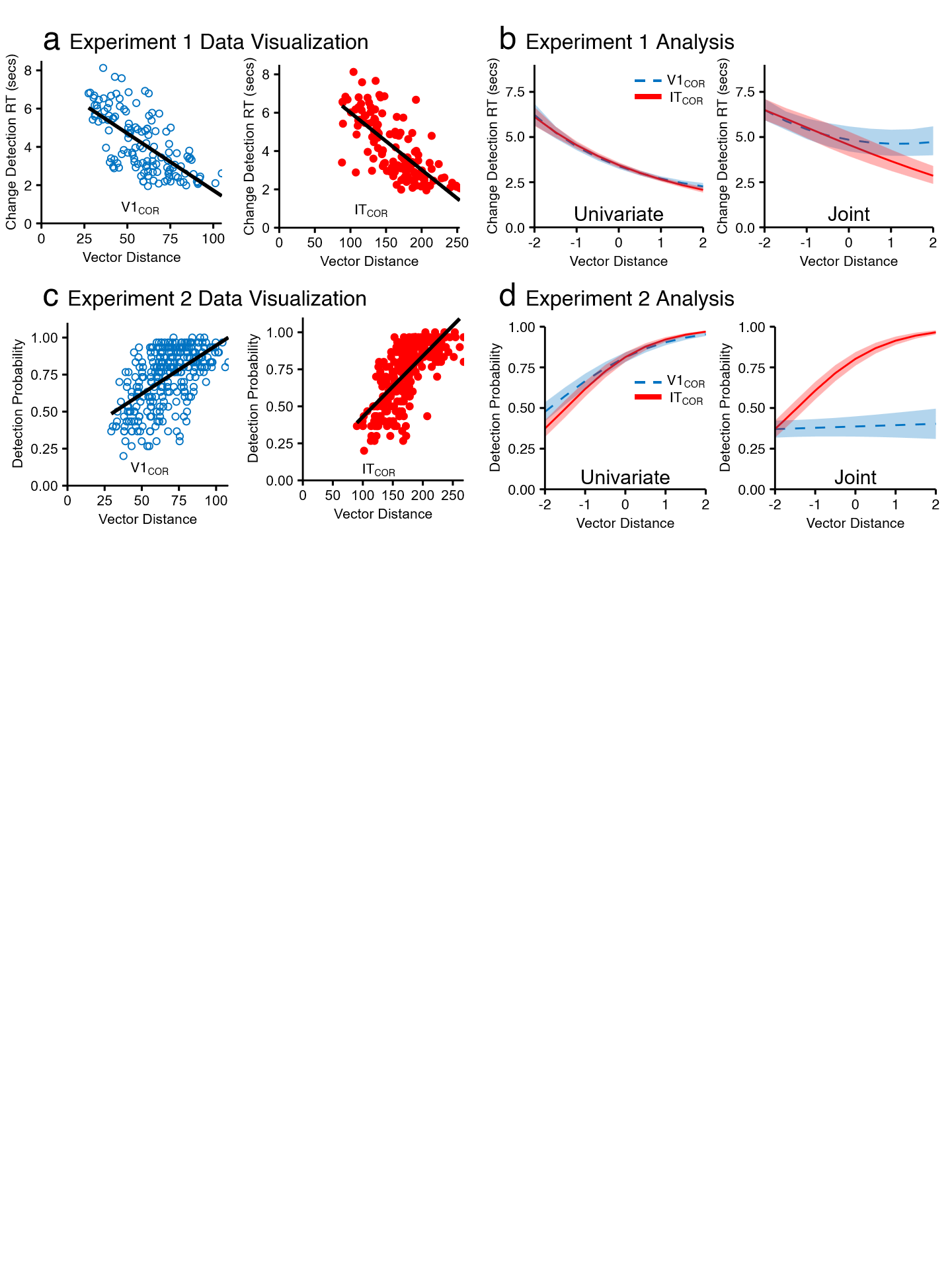

*Note.* **a**, Scatterplot visualization of the results from Experiment 1, showing the relationship between either the V1_COR_ or IT_COR_ vector distances for a given scene pair (without normalization) and the mean response time (RT) across participants for that pair. **c**, Scatterplot visualization of the results from Experiment 2, showing the relationship between either the V1_COR_ or IT_COR_ vector distances for a given scene pair (without normalization) and the proportion of participants who detected a change for that pair. A change was reported on 19.97% of no-change trials (not shown). **b & d**, Results of the statistical analysis of the single-trial RTs from Experiment 1 (**f**) and the single-trial binary responses from Experiment 2 (**d**). The univariate plots show predicted values when V1_COR_ and IT_COR_ were analyzed separately, whereas the joint plots shows predicted values when both variables were included simultaneously (with each effect being centered at -2 SD on the other variable). Shading indicates +/-1 SE.

**Figure S3**

*Experiment 3 results using unnormalized vectors.*
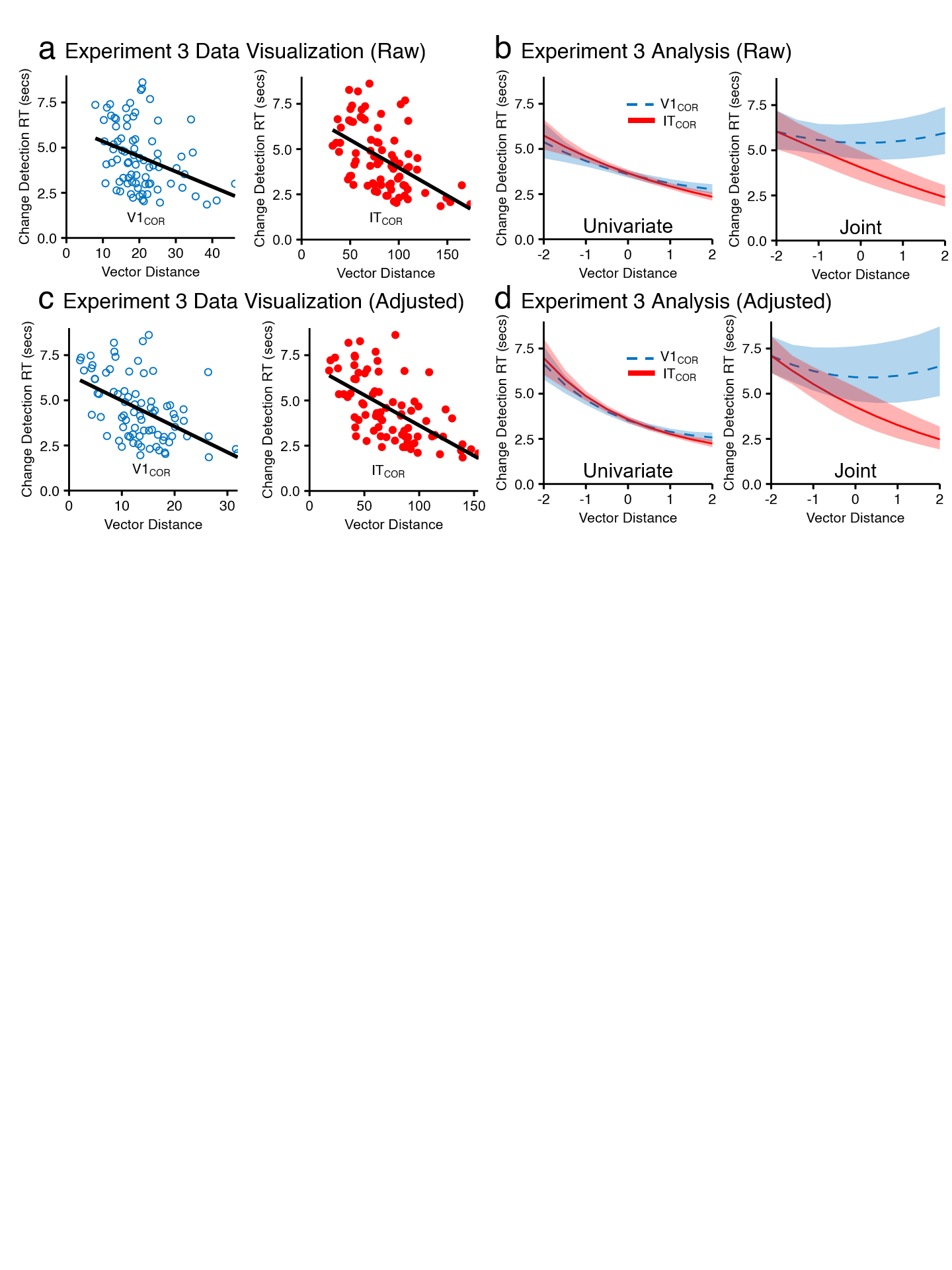

*Note.* **a**, Scatterplot visualization of the results from Experiment 3 with unnormalized vectors, showing the relationship between either the V1_COR_ or IT_COR_ vector distances for a given scene pair (without normalization) and the mean response time (RT) across participants for that pair. **b**, Results of the statistical analysis of the single-trial RTs from Experiment 3 with the unnormalized vector distances. Shading represents the standard error. The univariate plots show predicted values when V1_COR_ and IT_COR_ were analyzed separately, whereas the joint plot shows predicted values when both variables were included simultaneously (with each effect being centered at -2 SD on the other variable). **c, d**, Same as **a** and **b**, but using (unnormalized) vector distances computed after adjusting for the distance from the center of the image. Shading indicates +/- 1 SE.

**Figure S4**

*ERP waveforms*

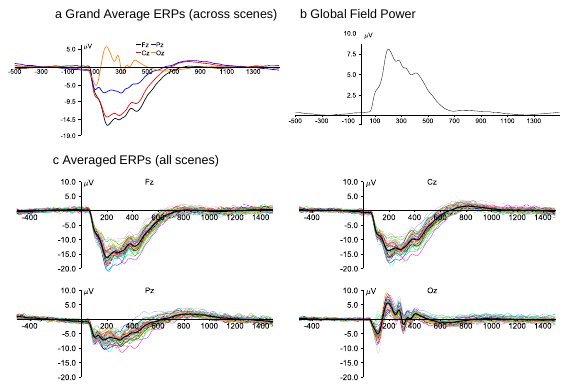

Note. **a**, Grand average event-related potentials (ERPs), collapsed across scenes and subjects, at four central electrode sites. **b**, Mean global field power collapsed across scenes and subjects. **c**, Averaged ERPs for each individual scene, collapsed across subjects at the four midline electrode sites. The grand average waveform is shown as the thick black line in each panel. Time zero is the onset of the scene in all waveforms.

**Figure S5**

*Representational Similarity Results for All 4 Areas of CORnet.*

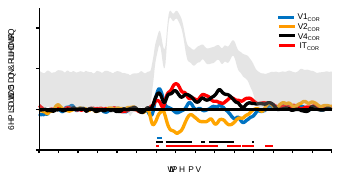

*Note.* Representational similarity between the V1_COR_, V2_COR_, V4_COR_, and IT_COR_ vectors and the pattern of voltage over the electrodes, computed separately at each time point for Experiment 4. This was done separately for each of the four areas, disregarding any shared variance. Horizontal line segments across the bottom indicated time periods in which the representational similarity values are significantly greater than zero (p <.05) after correction for multiple comparisons.

**Figure S6**

*Semipartial RSA correlations for Experiment 3*

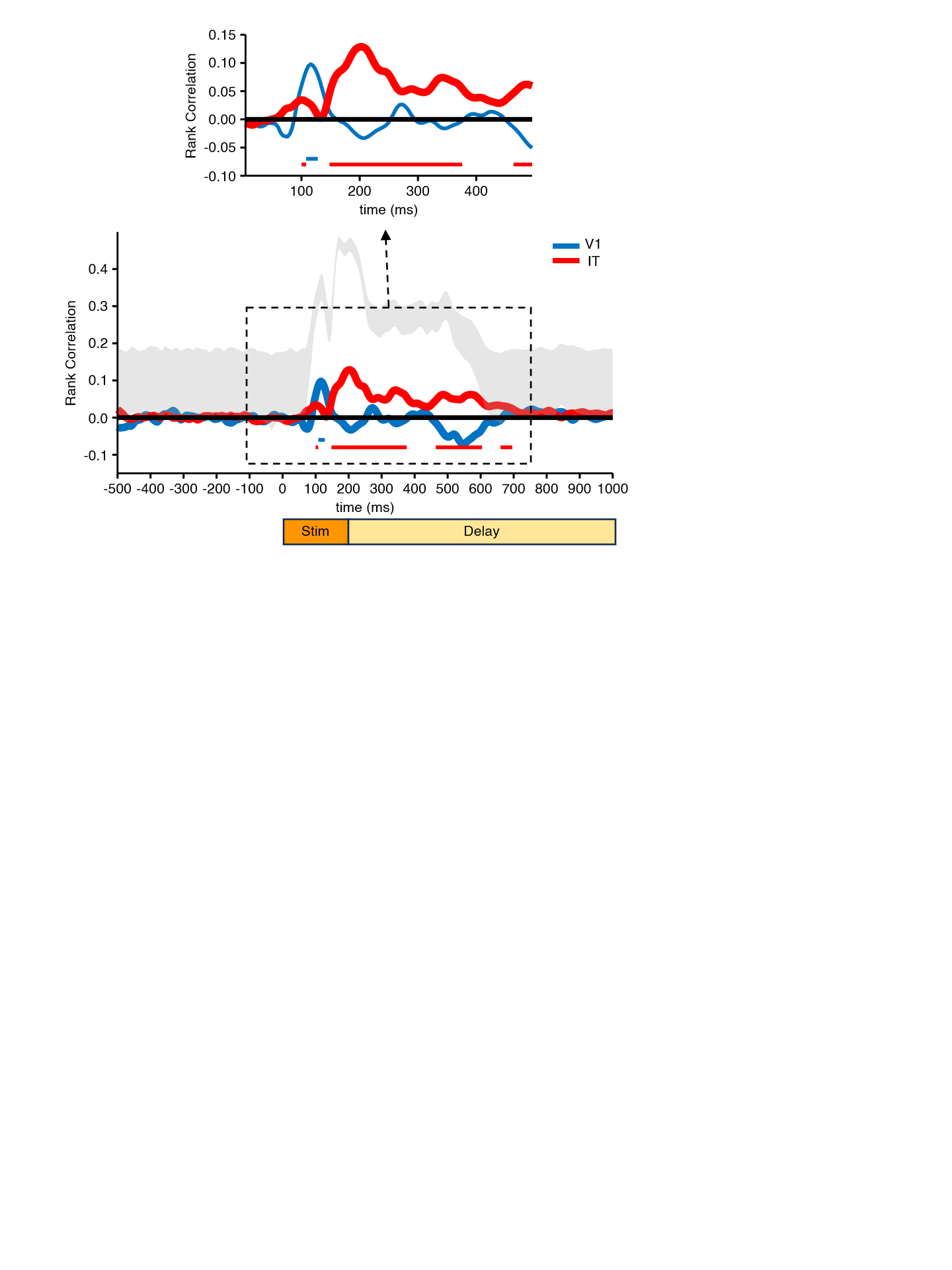

*Note.* Semipartial representational similarity correlations between the V1_COR_ or IT_COR_ vectors and the pattern of voltage over the electrodes, computed separately at each time point with the upper and lower bounds of the estimated noise ceiling of the ERP data shaded in gray. Horizontal line segments across the bottom indicated time periods in which the similarity values are significantly > 0 (FDR corrected p <.05).

**Table S1**

*Statistical results examining the unique contributions of V1_COR_ and IT_COR_ after controlling for similarity in Experiments 1 and 3*

|  | **Experiment 1** | **Experiment 3 (Adjusted)** | **Experiment 3 (Unadjusted)** |
| --- | --- | --- | --- |
| **V1****_COR_ + Similarity Model** | | | |
| Unique Contribution of Linear V1_COR_ | F(1,3719) = 13.11, p = .0003* | F(1,2455) = 18.24, p < .0001* | F(1,2455) = 2.17, p = .1405 |
| Unique Contribution of Linear Similarity | F(1,3719) = 181.56, p = .0001* | F(1,2455) = 138.84, p < .0001* | F(1,2455) = 144.22, p < .0001* |
| Unique Contribution of Quadratic V1_COR_ | F(1,3719) = 0.75, p = .3858 | F(1,2455) = 4.86, p = .0276* | F(1,2455) = 0.51, p = .4736 |
| Unique Contribution of Quadratic Similarity | F(1,3719) = 10.15, p = .0015* | F(1,2455) = 23.69, p < .0001 | F(1,2455) = 26.34, p < .0001 |
| **IT_COR_ + Similarity Model** | | | |
| Unique Contribution of Linear IT_COR_ | F(1,3719) = 8.68, p = .0032* | F(1,2455) = 23.99, p < .0001* | F(1,2455) = 2.08, p = .1494 |
| Unique Contribution of Linear Similarity | F(1,3719) = 114.61, p < .0001* | F(1,2455) = 98.46, p < .0001* | F(1,2455) = 122.38, p < .0001* |
| Unique Contribution of Quadratic IT_COR_ | F(1,3719) = .39, p = .5327 | F(1,2455) = 5.38, p = .0205* | F(1,2455) = 0.71, p = .3979 |
| Unique Contribution of Quadratic Similarity | F(1,3719) = 8.91, p = .0029* | F(1,2455) = 16.29, p < .0001* | F(1,2455) = 22.04, p < .0001* |

**Table S2**

*Locations (relative to the upper left corner) and sizes of the changed object in Experiment 3*

| Image Number | Center Of Mass Y Coordinate | Center Of Mass X Coordinate | Object Area In Total Pixels |
| --- | --- | --- | --- |
| 1 | 386.100699 | 250.938882 | 12026 |
| 2 | 150.019789 | 159.151715 | 3032 |
| 3 | 439.803313 | 249.329314 | 9538 |
| 4 | 368.035552 | 445.532527 | 3966 |
| 5 | 228.264933 | 415.543154 | 26116 |
| 6 | 291.949218 | 359.393947 | 3899 |
| 7 | 298.690681 | 134.101472 | 3262 |
| 8 | 130.213418 | 190.291491 | 17112 |
| 9 | 254.756619 | 312.010229 | 3324 |
| 10 | 405.402801 | 73.3005313 | 8282 |
| 11 | 369.778919 | 252.323063 | 22200 |
| 12 | 412.340069 | 452.696895 | 4058 |
| 13 | 334.907332 | 58.3790583 | 4651 |
| 14 | 411.103607 | 334.82079 | 20462 |
| 15 | 389.238609 | 304.539568 | 13344 |
| 16 | 341.413001 | 270.665349 | 4046 |
| 17 | 331.10261 | 297.454016 | 4980 |
| 18 | 431.036082 | 437.275629 | 7677 |
| 19 | 259.534656 | 284.127223 | 4386 |
| 20 | 298.827524 | 162.236747 | 2754 |
| 21 | 149.788034 | 291.082906 | 9360 |
| 22 | 266.917991 | 436.725251 | 8182 |
| 23 | 396.351553 | 415.647002 | 4153 |
| 24 | 452.339881 | 138.407143 | 8400 |
| 25 | 311.168126 | 78.5597632 | 8107 |
| 26 | 228.80129 | 122.202703 | 3256 |
| 27 | 86.5879206 | 254.401893 | 9719 |
| 28 | 297.729763 | 461.725885 | 4126 |
| 29 | 321.667134 | 386.33876 | 3563 |
| 30 | 113.64267 | 298.088112 | 11474 |
| 31 | 459.306723 | 399.27395 | 3570 |
| 32 | 413.945829 | 293.299025 | 2769 |
| 33 | 301.703149 | 431.308209 | 1937 |
| 34 | 281.780312 | 386.027011 | 4998 |
| 35 | 402.497302 | 443.725045 | 4448 |
| 36 | 449.548258 | 389.140523 | 5999 |
| 37 | 389.229174 | 371.012654 | 5690 |
| 38 | 432.653298 | 454.011991 | 4837 |
| 39 | 358.694564 | 275.9035 | 6715 |
| 40 | 289.719098 | 252.342706 | 7540 |
| 41 | 311.353448 | 109.687979 | 4176 |
| 42 | 306.486338 | 91.5291287 | 5819 |
| 43 | 432.147122 | 143.368159 | 1407 |
| 44 | 307.546043 | 196.005036 | 5560 |
| 45 | 454.076679 | 84.0029085 | 3782 |
| 46 | 355.523793 | 399.434262 | 5674 |
| 47 | 425.860373 | 293.766106 | 15212 |
| 48 | 293.119169 | 237.202771 | 4330 |
| 49 | 172.009677 | 354.78919 | 7234 |
| 50 | 191.451566 | 146.578963 | 10220 |
| 51 | 209.506688 | 364.192312 | 8299 |
| 52 | 372.171158 | 366.136693 | 11171 |
| 53 | 300.124151 | 106.911283 | 63922 |
| 54 | 62.4186121 | 283.576783 | 9338 |
| 55 | 315.078209 | 435.049938 | 9692 |
| 56 | 342.626412 | 164.260859 | 3983 |
| 57 | 435.48097 | 387.29536 | 3836 |
| 58 | 280.908131 | 303.373925 | 5116 |
| 59 | 131.558861 | 414.660221 | 14118 |
| 60 | 141.104467 | 281.119473 | 8730 |
| 61 | 159.288535 | 197.603779 | 14078 |
| 62 | 272.809011 | 145.302069 | 3529 |
| 63 | 304.422927 | 232.553182 | 4353 |
| 64 | 237.936968 | 264.111882 | 3173 |
| 65 | 275.977483 | 399.683554 | 14034 |
| 66 | 205.524254 | 416.748134 | 2144 |
| 67 | 169.259993 | 162.257704 | 22716 |
| 68 | 199.847484 | 422.893472 | 21506 |
| 69 | 258.716945 | 172.851412 | 3116 |
| 70 | 395.347888 | 106.828987 | 4947 |
| 71 | 232.766483 | 210.378644 | 31972 |
| 72 | 370.033103 | 53.5281748 | 12174 |
| 73 | 121.668071 | 319.009978 | 9020 |
| 74 | 187.08254 | 302.219487 | 9668 |
| 75 | 448.804416 | 351.081668 | 2853 |
| 76 | 211.768687 | 81.0869975 | 6368 |
| 77 | 411.712921 | 412.199211 | 4814 |
| 78 | 444.035758 | 182.663808 | 8977 |
| 79 | 325.382631 | 31.7851306 | 6241 |
| 80 | 379.863304 | 259.669447 | 4916 |

**Table S3**

*Statistical results for V2_COR_ and V4_COR_ in Experiments 1–3*

|  | **Experiment 1** | **Experiment 2** | **Experiment 3 (Adjusted)** | **Experiment 3 (Unadjusted)** |
| --- | --- | --- | --- | --- |
| **V2****_COR_ Only Models** | r^2^ = .707 | r^2^ = .624 | r^2^ = .391 | r^2^ = .284 |
| Unique Contribution of Linear V2_COR_ | F(1,3721) = 266.41,  p < .0001* | F(1,8854) = 403.95,  p < .0001* | F(1,2457) = 37.35,  p < .0001* | F(1,2457) = 17.68,  p < .0001* |
| Unique Contribution of Quadratic V2_COR_ | F(1,3721) = 0.02,  p = .8763 |  | F(1,2457) = 1.26,  p = .2622* | F(1,2457) = .92,  p = .9191 |
| **V4_COR_ Only Models** | r^2^ = .726 | r^2^ = .672 | r^2^ = .423 | r^2^ = .327 |
| Unique Contribution of Linear V4_COR_ | F(1,3721) = 287.74,  p < .0001* | F(1, 8854) = 481.78,  p < .0001* | F(1,2457) = 41.07,  p < .0001* | F(1,2457) = 20.86,  p < .0001* |
| Unique Contribution of Quadratic V4_COR_ | F(1,3721) = 1.02,  p = .3121 |  | F(1,2457) = 0.34,  p = .5587 | F(1,2457) = .14,  p = .7100 |

**Table S4**

*Test statistics for analyses of single-trial response times in Experiment 1 with Unnormalized Distances*

|  | **Experiment 1** |
| --- | --- |
| Unique Contribution of Linear V1_COR_ | F(1,3721) = 110.06, p < .0001* |
| Unique Contribution of Quadratic V1_COR_ | F(1,3721) = 1.53, p = .2158 |
| Unique Contribution of Linear IT_COR_ | F(1,3721) = 150.94, p < .0001* |
| Unique Contribution of Quadratic IT_COR_ | F(1,3721) = 0.20, p = .6550 |
| Unique Contribution of Linear V1_COR_ | F(1,3719) = 3.85, p = .0498* |
| Unique Contribution of Linear IT_COR_ | F(1,3719) = 26.16, p < .0001* |
| Unique Contribution of Quadratic V1_COR_ | F(1,3719) = 1.50, p = .2204 |
| Unique Contribution of Quadratic IT_COR_ | F(1,3719) = 0.32, p = .5707 |

**Table S5**

*Test statistics for analyses of single-trial accuracy (Experiment 2) with Unnormalized Distances*

| **V1****_COR_ Only Analysis** | |
| --- | --- |
| Unique Contribution of Linear V1_COR_ | F(1,8854) = 138.39, p < .0001* |
| **IT_COR_ Only Analysis** | |
| Unique Contribution of Linear IT_COR_ | F(1,8854) = 288.22, p < .0001* |
| **V1_COR_ + IT_COR_ Analysis** | |
| Unique Contribution of Linear V1_COR_ | F(1,8853) = 0.15, p = 0.6967 |
| Unique Contribution of Linear IT_COR_ | F(1,8853) = 111.02, p < .0001* |

**Table S6**

*Test statistics for analyses of single-trial response times (Experiments 3) with Unnormalized Distances*

|  | | **Experiment 3 (Adjusted)** | **Experiment 3 (Unadjusted)** |
| --- | --- | --- | --- |
| Unique Contribution of Linear V1_COR_ | | F(1,2457) = 33.91, p < .0001* | F(1,2457) = 9.41, p = .0022* |
| Unique Contribution of Quadratic V1_COR_ | | F(1,2457) = 2.39, p = .1223 | F(1,2457) = 0.35, p = .5561 |
| Unique Contribution of Linear IT_COR_ | | F(1,2457) = 57.65, p < .0001* | F(1,2457) = 28.35, p < .0001* |
| Unique Contribution of Quadratic IT_COR_ | | F(1,2457) = 0.77, p = 0.3808 | F(1,2457) < 0.01, p = .9656 |
| Unique Contribution of Linear Similarity | | F(1,2457) = 175.75, p < .0001* | |
| Unique Contribution of Quadratic Similarity | | F(1,2457) = 29.56, p < .0001* | |
| Unique Contribution of Linear V1_COR_ | | F(1,2455) = 0.10, p = .7462 | F(1,2455) < 0.01, p = .9562 |
| Unique Contribution of Linear IT_COR_ | | F(1,2455) = 15.08, p = .0001* | F(1,2455) = 18.10, p < .001* |
| Unique Contribution of Quadratic V1_COR_ | | F(1,2455) = 1.20, p = .2741 | F(1,2455) = 0.71, p = .3995 |
| Unique Contribution of Quadratic IT_COR_ | | F(1,2455) = 0.01, p = .9053 | F(1,2455) = 0.24, p = .6248 |
| Unique Contribution of Adjusted Linear V1_COR_ | |  | F(1,2455) = 18.27, p < .0001* |
| Unique Contribution of Unadjusted Linear V1_COR_ | |  | F(1,2455) = 0.79, p = 0.3739 |
| Unique Contribution of Quadratic Adjusted V1_COR_ | |  | F(1,2455) = 1.28, p = 0.2578 |
| Unique Contribution of Quadratic Unadjusted V1_COR_ | |  | F(1,2455) = 0.11, p = .7417 |
| Unique Contribution of Adjusted Linear IT_COR_ | |  | F(1,2455) = 21.98, p < .0001* |
| Unique Contribution of Unadjusted Linear IT_COR_ | |  | F(1,2455) = 0.24, p = .6238 |
| Unique Contribution of Quadratic Adjusted IT_COR_ | |  | F(1,2455) = 0.33, p = .5668 |
| Unique Contribution of Quadratic Unadjusted IT_COR_ | |  | F(1,2455) = 1.24, p = .2657 |

**Table S7**

*Unnormalized Vector Length Means and Standard Deviations <- add normalized and mention in new section?*

|  | **Experiment 1** | **Experiment 2** | **Experiment 3 (Adjusted)** | **Experiment 3 (Unadjusted)** |
| --- | --- | --- | --- | --- |
| V1 | 58.97 (17.80) | 67.39 (17.10) | 13.45 (6.34) | 20.23 (7.40) |
| V2 | 98.66 (21.90) | 109.75 (19.45) | 27.93 (9.90) | 31.50 (10.20) |
| V4 | 56.17 (13.47) | 62.11 (12.00) | 20.73 (8.00) | 22.72 (7.98) |
| IT | 161.42 (38.48) | 174.13 (33.48) | 74.05 (33.28) | 83.01 (29.95) |
